# Soil pH as an external filter shaping insect–microbe gut symbiosis

**DOI:** 10.1101/2025.11.17.688959

**Authors:** Hideomi Itoh, Hiroyuki Shimoji, Daisuke Nakane, Seonghan Jang, Yoshitomo Kikuchi

## Abstract

**Background:** Many animals and plants establish intimate symbiotic relationships with specific microorganisms acquired from the environment. Given the immense diversity of environmental microbiomes, selecting appropriate partners from such a vast microbial pool poses a critical challenge for host organisms. To meet this challenge, hosts have evolved sophisticated internal partner-choice mechanisms that ensure stable associations with beneficial microbes. However, because these symbionts primarily inhabit external environments, environmental conditions themselves are also expected to play pivotal roles in the establishment of symbiosis. Despite this expectation, the mechanistic role of external environmental filters in shaping the intended symbiosis remains largely unexplored. Focusing on stink bugs, which acquire their symbiotic bacteria from soil each generation, we investigated how soil properties influence the establishment of gut symbiosis in terrestrial insects.

**Results:** Microbiome analyses confirmed that *Burkholderia sensu lato* overwhelmingly dominates a specific gut organ in six stink bug species from the superfamilies Coreoidea and Lygaeoidea, including serious agricultural pests (relative abundance ranging from 74.5% to 100%). Rearing experiments with isolated *Burkholderia* revealed that insects were strictly dependent on this symbiont; failure to acquire it from soil severely reduced host growth and reproduction, indicating that the availability of symbionts from soil represents an ecological bottleneck. Field surveys identified patches of exceptionally high stink bug density in weedy fields with soil pH <7.0, whereas such aggregations were absent in fields with pH ≥7.0. Laboratory experiments with collected field soils showed that the abundance of *Burkholderia* in soils was negatively correlated with soil pH, and stink bugs readily acquired their symbionts from soils with pH <7.0 but rarely from soils with pH ≥7.0. Experimental manipulations of soil pH followed by rearing experiments confirmed that increasing soil pH to 7–8 markedly suppressed symbiont acquisition by the host, likely by impairing symbiont growth and motility.

**Conclusions:** We demonstrate that, beyond host-intrinsic mechanisms, a soil chemical property externally filters symbiotic bacteria prior to colonization inside the host. These findings reveal that, alongside host physiology, environmental conditions—mediated through the soil and host gut microbiomes—shape the assembly, maintenance, and evolution of host–microbe symbiotic associations.

## Introduction

Symbiosis with microorganisms is ubiquitous among animals and plants and has been a major driver of organismal evolution [1]. Given the enormous diversity of microbes in the environment, establishing and maintaining mutualistic relationships with specific partners is a considerable challenge for host organisms. Many insects and some marine invertebrates have evolved vertical transmission systems to ensure partner fidelity [2–4]. By contrast, numerous animals and plants do not vertically transmit their symbionts but instead acquire them from the environment every generation. To secure partner specificity under these conditions, hosts have evolved sophisticated partner choice mechanisms. For example, in the legume–*Rhizobium* nodule symbiosis and the squid–*Vibrio* bioluminescent symbiosis, strict partner selection is mediated through chemical crosstalk between host and symbiont [5, 6]. In many animals, the composition of gut microbial communities is shaped by gut physiology and immunity [7].

In environmentally acquired symbioses, the environment has often been regarded simply as a reservoir of symbiotic microorganisms. However, microbial distributions in the environment are far from uniform. Global-scale surveys have revealed pronounced geographic variation in bacterial and fungal communities [8, 9], while more localized studies show substantial heterogeneity even among nearby sites [10]. From this perspective, the uneven distribution of symbiotic microorganisms in the environment may directly influence the establishment and evolution of symbioses. This process can be regarded as external, or pre-infection, selection of symbiotic microorganisms. Despite its potential importance, meanwhile, the role of environmental heterogeneity in shaping host–microbe specificity remains largely unexplored.

Phytophagous species of the superfamilies Coreoidea and Lygaeoidea of the order Heteroptera commonly establish gut symbioses with bacteria in *Burkholderia sensu lato*, including the genera *Caballeronia* and *Paraburkholderia* [11–18]. These symbionts are acquired from ambient soil during the early larval stage in every host generation and are specifically harbored in a crypt-bearing region of the posterior midgut [11, 19], where they enhance host survival and growth [11, 12] by performing pivotal metabolic roles [20, 21]. The multilayered host-internal mechanisms of symbiont filtering from environment microbiota have been well investigated in the bean bug *Riptortus pedestris* and its symbiont *Caballeronia insecticola*. The midgut of *R. pedestris* is divided into regions associated with food digestion (M1, M2, and M3) and the symbiotic organ (M4) [22]. Between the M3 and M4 regions lies a symbiont-sorting organ, the constricted region (CR), characterized by an extremely narrow lumen of only a few micrometers in diameter [22]. The CR, filled with a mucus-like matrix, functions as a physical barrier, preventing most non-symbiotic bacteria from passing through [22]. *Caballeronia* symbionts, in contrast, can traverse the CR by utilizing a unique flagellar-wrapping motility, allowing them to successfully colonize the symbiotic organ [22, 23]. In addition, the M3 and M4 regions exhibit high levels of digestive enzymes and antimicrobial peptides, which play crucial roles in selecting the symbiotic bacteria [24]. Furthermore, the CR gate is permanently sealed after symbiont passage and colonization in M4, thereby preventing secondary inoculation of contaminants [25].

Here we show that, beyond these host-internal filters such as physical barriers and antimicrobial peptides [22, 24, 25], an overlooked but critical layer of symbiont selection occurs externally, in the soil. We first identified the reliance on soil-derived *Burkholderia* in six stink bug species within the superfamilies Coreoidea and Lygaeoidea, including serious rice pests whose symbiotic associations have remained poorly characterized. Then, through characterization, field survey, and rearing experiments, we discovered that soil pH strongly regulates the density and activity of the symbiotic bacteria, and that symbiosis cannot be established in soils with pH ≥ 7. This observed dependence of gut symbiosis on soil fundamentally broadens current understanding of host–microbe interactions by demonstrating that external environmental properties, not only host-internal physiology, can dictate the establishment and evolution of gut symbioses.

## Results

### Obligate symbiosis with soil bacteria

Wild populations of the stink bug species examined in this study, *Cletus punctiger*, *C. schmidti*, *R. pedestris*, *Leptocorisa chinensis*, *Pachygrontha antennata*, and *Metochus uniguttatus*, possess cryptic or tubular organs that develop at the posterior region of the midgut (symbiotic organ: M4B and M4 in Fig. 1A–C and Fig. S1), where the bacterial symbionts colonize [26]. Microbiome analysis of field-captured adult insects showed that 97.6%–100% of individuals were positive for *Burkholderia* (Fig. 1D and Table S1), which consistently dominated the symbiotic organ, with relative abundance ranging from 74.5% to 100% (Fig. S2). Eggs laid by field-captured females and insects reared in clean Petri dishes were negative for *Burkholderia* (Fig. 1E and Table S1). Similarly, except for *C. schmidti*, all reared insects with an adult pair were also negative for *Burkholderia* (Table S1). Conversely, 93.8%–100% of insects reared with the soil have acquired *Burkholderia* (Fig. 1F and Table S1). In the rearing experiments conducted in the plant pots, all insect individuals reared under soil-accessible conditions were infected with *Burkholderia*, whereas those isolated from the soil and accessible only to the plant were not (Fig. S3A–C). These results revealed that these stink bug species are entirely dependent on gut symbionts existing in the soil, although *C. schmidti* has shown partial vertical transmission.

**Figure 1.**
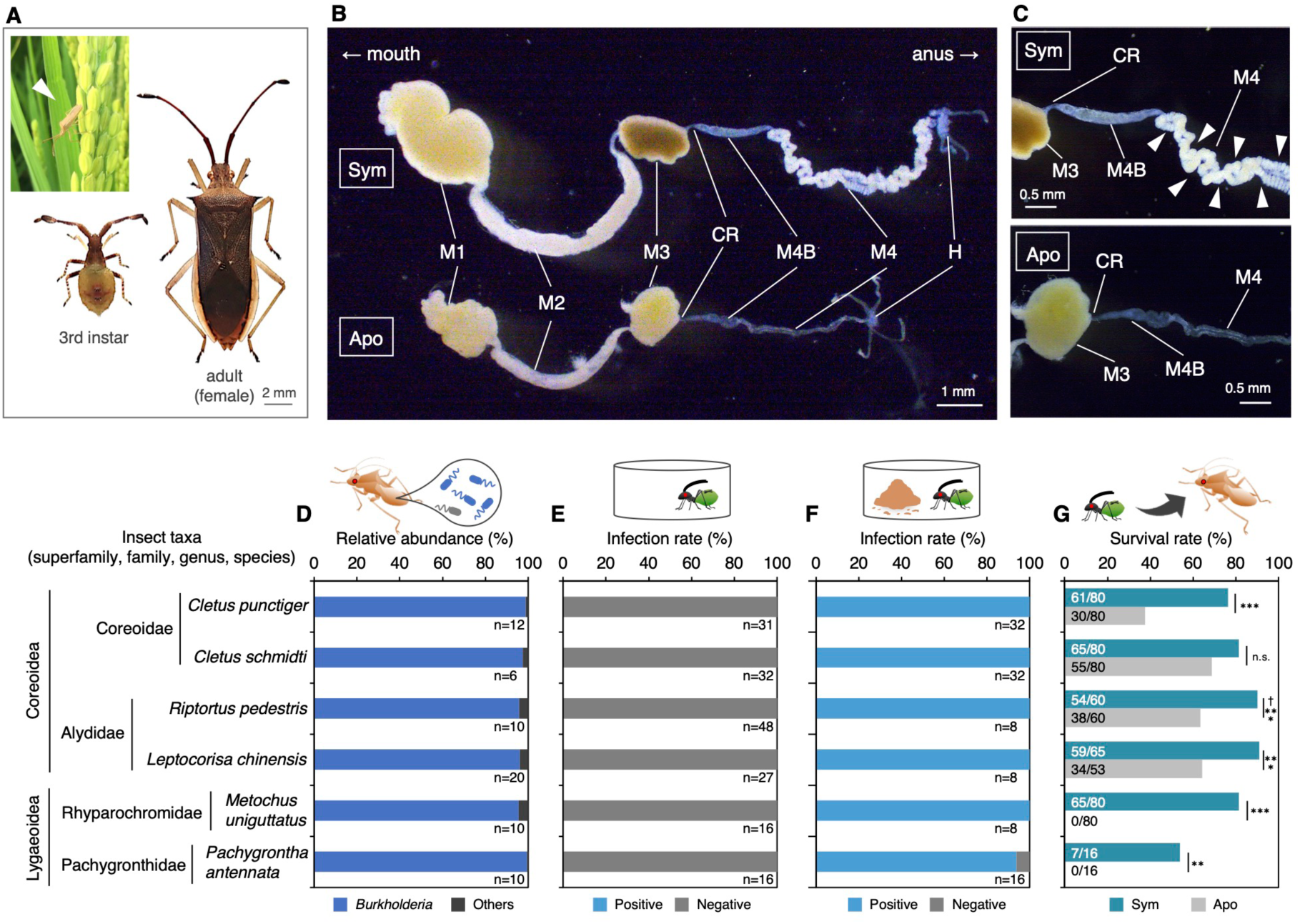
Characterization of gut symbiosis in the stink bugs used in this study. (A) The third instar and adult female of laboratory-maintained *C. punctiger*. The inset displays a wild individual of *C. punctiger* invading a rice paddy field (closed triangle). (B) Each dissected whole gut of the 4th instar of *C. punctiger* infected with *Burkholderia* (Sym) or uninfected (Apo). Abbreviations: M1, midgut first section; M2, midgut second section; M3, midgut third section; CR, constricted region; M4B, midgut fourth section with bulb; M4, midgut fourth section with crypts (symbiotic organ); H, hindgut. (C) Magnified photo of the M3 to M4 region of Sym and Apo insects. The closed triangles in M4 indicate crypts. Photos of other stink bug species and their whole guts used in this study are shown in Fig. S1. (D) Community structure of the gut microbiota in wild populations of six stink bug species. Data represent the average composition, with the number of examined individuals shown below each bar. Detailed sample information and individual gut microbiota community structures are provided in Table S7 and Fig. S2, respectively. (E) and (F) Infection rates to *Burkholderia* in insects reared without (E) and with soil (F). Values below each bar indicate the number of examined individuals. (G) Survival rates (i.e., adult emergence rates) of six stink bug species infected with *Burkholderia* (Sym) or uninfected (Apo). ^†^Data of *R. pedestris* are from a previous study [27]. Numbers on the bars represent “number of survivors/total number of examined individuals”. Statistical analysis was performed using Fisher’s exact test (**, *P* < 0.01; ***, *P* < 0.001; n.s., not significant (*P* ≥ 0.05)).

In the absence of soil-derived *Burkholderia* symbionts, survival rates were significantly reduced in all species except *C. schmidti* (Fig. 1G). Notably, *M. uniguttatus* and *P. antennata* failed to reach adulthood without acquiring *Burkholderia* (Fig. 1G). Although aposymbiotic individuals of *Cletus* species were not entirely incapable of reaching adulthood, they exhibited multiple morphological and reproductive abnormalities. Specifically, aposymbiotic *Cletus* individuals exhibited a decline in growth rate, dwarfed body size, and forewing atrophy (Fig. 2A–E). Furthermore, aposymbiotic adults suffered reproductive impairments, including immature reproductive organs, reduced mating behavior, lower egg production, and decreased hatching rates (Fig. 2F–J). These results demonstrate that the tested stink bug species are strictly dependent on soil-borne *Burkholderia*, indicating that the availability of *Burkholderia* from the ambient soil is a critical factor for development, reproduction, and population maintenance.

**Figure 2.**
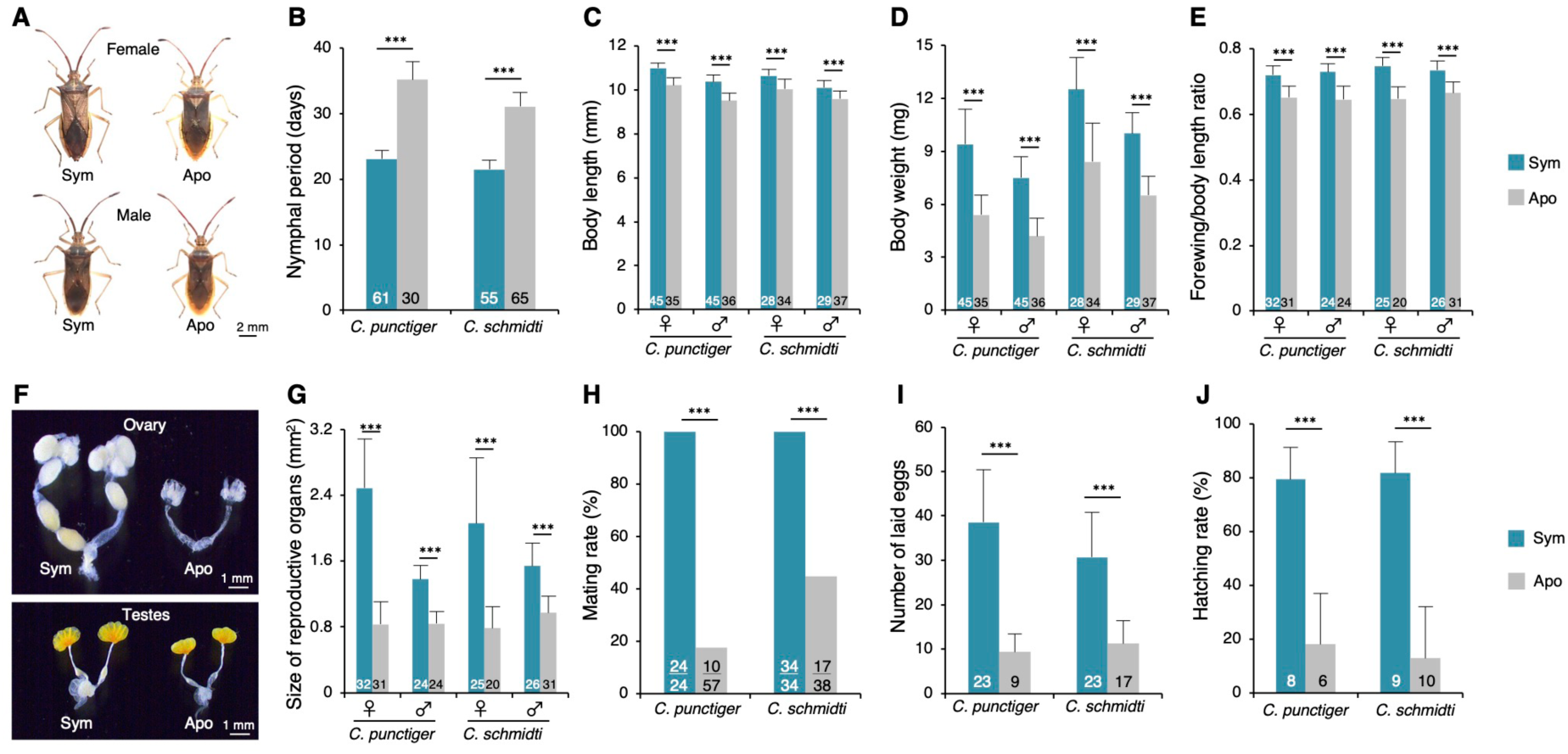
Effects of *Burkholderia* symbiosis on host growth and fecundity in *Cletus* spp. Fitness effects of *Burkholderia* symbionts on insect growth (A–E) and fecundity (F–J). (A) Sym and Apo adult females (upper) and male (lower) of *C. punctiger*. (B) Developmental period (days) from hatching to adulthood. (C) Adult body length. (D) Whole-body dry weight. (E) Ratio of forewing length to body length in adult insects. (F) Reproductive organs in Sym and Apo adults: ovary (upper) and testes (lower). (G) Size of reproductive organs. (H) Mating rate. (I) Total number of eggs laid within two weeks after first oviposition. (J) Hatching rate of eggs laid by Sym and Apo females. Numbers in bars indicate “total number of examined insect samples” (B–E, G, I–J) or “number of successful pairs/total number of examined pairs” (H). Error bars represent standard deviation. Statistical analyses were performed using Student’s t-test, Welch’s t-test, Mann–Whitney *U* test, or Brunner–Munzel test after checking for normality and homogeneity of variance (***, *P* < 0.001) (B–E, G, I–J), or using Student’s t-test (***, *P* < 0.001) (H).

### Symbiosis establishment dependent on soil pH

A number of studies have investigated the environmental determinants of soil microbiome diversity, consistently identifying soil pH as one of the most influential physicochemical parameters shaping microbial community structure and diversity; it exerts a substantial impact on microbial assemblages, often overriding other edaphic factors such as soil type, moisture, or nutrient content [8, 28–33]. Notably, a continent-wide survey reported that *Burkholderia sensu lato* are more prevalent in mildly acidic soils than in neutral or alkaline conditions [34]. Collectively, these findings suggest that soil pH may affect the local distribution of symbiotic *Burkholderia* and, by extension, influence the ecological interactions of stink bugs that rely on the environmental acquisition of these symbionts. To clarify this hypothesis, we surveyed stink bug biomass across weedy fields with varying soil pH (Fig. 3A and Table S2). The major Coreoidea and Lygaeoidea species captured in these field surveys were *C. punctiger*, *L. chinensis*, and *Paromius jejunus* (Coreoidea) (Fig. S4A). Although the size of the stink bug population was not significantly different among fields, large numbers of nymphs and adults were observed at several sites with soil pH <7.0 (Fig. 3B). Sensitivity analyses confirmed that these high-abundance data points were outliers (Fig. S5B), suggesting that localized stink bug surges tend to occur in mildly acidic soil environments.

**Figure 3.**
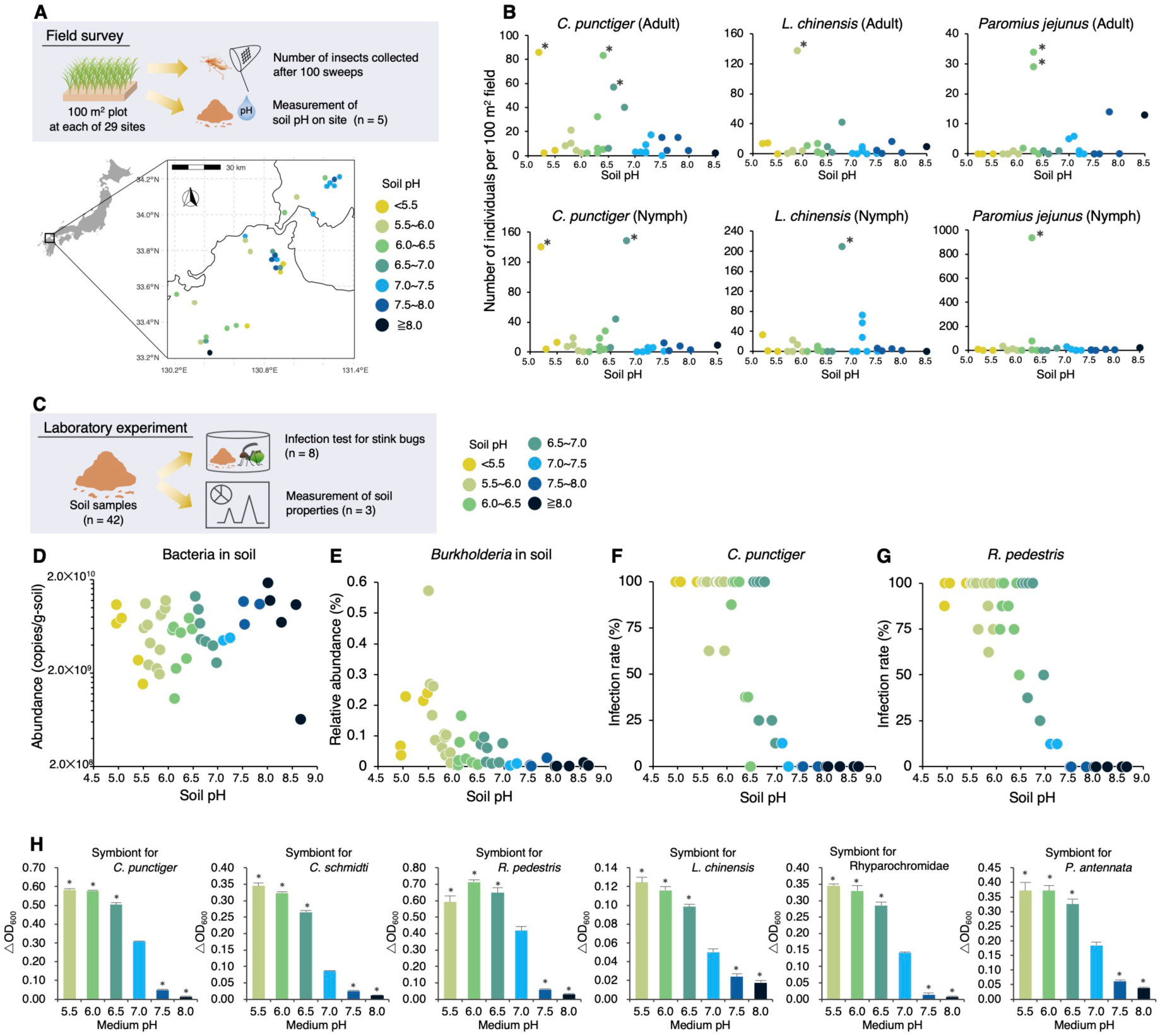
Soil pH preferences for symbiosis establishment inferred from field surveys and laboratory experiments. (A) Schematic overview of the field survey and geographic distribution of the examined weedy fields. Detailed site information and results are provided in Table S2. (B) Number of stink bugs collected from each weedy field for each insect species. The left and right panels show the number of adults and nymphs, respectively. Data points marked with an asterisk indicate outbreaks identified through statistical analysis (Fig. S4B). Colors represent variations in soil pH. (C) Schematic diagram of the laboratory infection test conducted on insects using soil samples. Detailed information of the soils is summarized in Table S3. (D) Bacterial abundance in the examined soil samples, estimated using qPCR analysis. (E) Relative abundance of *Burkholderia* in each soil microbiome. (F) and (G) Correlation between soil pH and infection rates of *Burkholderia* in *C. punctiger* (F) and *R. pedestris* (G), respectively. Each data point represents the infection rate among 8 individuals. (H) pH preferences for the growth of symbiotic *Burkholderia* strains derived from stink bug species. Error bars represent standard deviation (n = 3). Values marked with an asterisk indicate a significant difference compared with those at pH 7.0 (*, *P* < 0.05), as determined by Dunnett’s test.

On the other hand, because field surveys inevitably involve several uncontrolled factors—such as insecticide use, differences in weeding schedules, and variation in plant biomass—these local environmental variations may have obscured or weakened the apparent relationship between soil conditions and symbiosis establishment. To better evaluate the specific impact of soil pH under controlled conditions, we next conducted laboratory experiments. Using soil samples collected from agricultural and weedy fields across Japan (Table S3), we analyzed the relationships among soil pH and *Burkholderia* availability of stink bugs (Fig. 3C). Soil pH did not correlate with the total bacterial density (Generalized Linear Model (GLM) with Gaussian, LR test, *χ*^2^ test, *χ*^2^ = 1.580, *df* = 1, *P* = 0.205, *R*^2^ = 0.075; Fig. 3D and Table S4), however it significantly negatively correlated with the relative abundance of *Burkholderia* (GLM with beta-binomial, LR test, *χ*^2^ = 23.589, *df* = 1, *P* < 0.001, *R*^2^ = 0.016, *Q* < 0.001; Fig. 3E and Table S4). Among 14 chemical variables, soil pH and exchangeable calcium (which is sensitive to soil pH) were identified as the main factors influencing the relative abundance of *Burkholderia* in the soil microbiota (Table S5), consistent with a previous continental-scale study reporting a negative correlation between soil pH and *Burkholderia* abundance [34]. Afterward, to assess whether soil pH affects symbiont acquisition, the collected soils were fed with nymphs of *C. punctiger* and *R. pedestris*. The infection rates of both stink bug species drastically declined in soils with pH ≥7.0, whereas soils with pH <7.0 supported efficient symbiont acquisition (*C. punctiger*: GLM with beta-binomial, LR test, *χ*^2^ = 43.857, *df* = 1, *P* < 0.001, *R*^2^ = 0.366; *R. pedestris*: GLM with beta-binomial, LR test, *χ*^2^ = 33.09, *df* = 1, *P* < 0.001, *R*^2^ = 0.312; Fig. 3F–G and Table S4). This pH-dependence of *Burkholderia* availability was also observed in four other stink bug species (Fig. S5A). These results strongly suggest that soil pH strongly affects symbiont acquisition by stink bugs, with pH 7.0 representing as a tipping point for the success or failure of symbiont acquisition in the field.

To confirm how pH affects symbiont growth, we examined the *in vitro* growth rates of symbiotic *Burkholderia* strains derived from stink bugs, along with some type strains within the *Burkholderia sensu lato*, under different pH levels. Our results demonstrated that all *Burkholderia* strains grew well under low pH conditions (<7.0) compared to pH 7.0 (Fig. 3H and Fig. S5B). However, under high pH conditions (>7.0), the growth rate of *Burkholderia* significantly decreased (Fig. 3H and Fig. S5B). Since *Burkholderia* symbionts positively affected host fitness; particularly concerning survival, growth, and fecundity (Figs. 1G and 2), these results suggest that mildly acidic soils provide a favorable environment for stink bugs by enhancing the availability of beneficial symbionts that improve host fitness.

### Experimental modification of soil pH inhibits the establishment of symbiosis

To further investigate the causal relationship between soil pH and symbiosis establishment, we experimentally increased the pH of mildly acidic soils with pH 5.6 (pH 5.6 fraction) and 5.9 (pH 5.9 fraction) using CaCO_3_ (Fig. 4A–B). In these pH-adjusted soils (maintained at pH 7–8), the infection rate of *C. punctiger* significantly decreased as soil pH increased in both mildly acidic soils (pH 5.6 fraction: GLM with beta-binomial, LR test, *χ*^2^ = 45.968, *df* = 1, *P* < 0.001, *R*^2^ = 0.248; pH 5.9 fraction: GLM with beta-binomial, LR test, *χ*^2^ = 25.943, *df* = 1, *P* < 0.001, *R*^2^ = 0.261; Fig. 4C and Table S4). A similar negative effect of elevated pH on symbiosis was observed for *R. pedestris* (pH 5.6 fraction: GLM with beta-binomial, LR test, *χ*^2^ = 46.17, *df* = 1, *P* < 0.001, *R*^2^ = 0.304; pH 5.9 fraction: GLM with beta-binomial, LR test, *χ*^2^ = 34.715, *df* = 1, *P* < 0.001, *R*^2^ = 0.228; Fig. 4D and Table S4). In parallel, *Burkholderia* abundance in soil also declined with increasing soil pH (pH 5.6 fraction: GLM with Gaussian, LR test, *χ*^2^ = 10.237, *df* = 1, *P* < 0.001, *R*^2^ = 0.145; pH 5.9 fraction: GLM with Gaussian, LR test, *χ*^2^ = 20.736, *df* = 1, *P* < 0.001, *R*^2^ = 0.385; Fig. 4E and Table S4). Similarly, negative impacts were detected on infection rates when soil pH was increased using steel slag or biochar instead of CaCO_3_ (Fig. 4F). The overall results from our laboratory experiments, in conjunction with the and field surveys, demonstrated that soil pH of ≥7.0 inhibits symbiosis establishment via the reduced abundance of *Burkholderia* in the soil, thereby regulating insect biomass.

**Figure 4.**
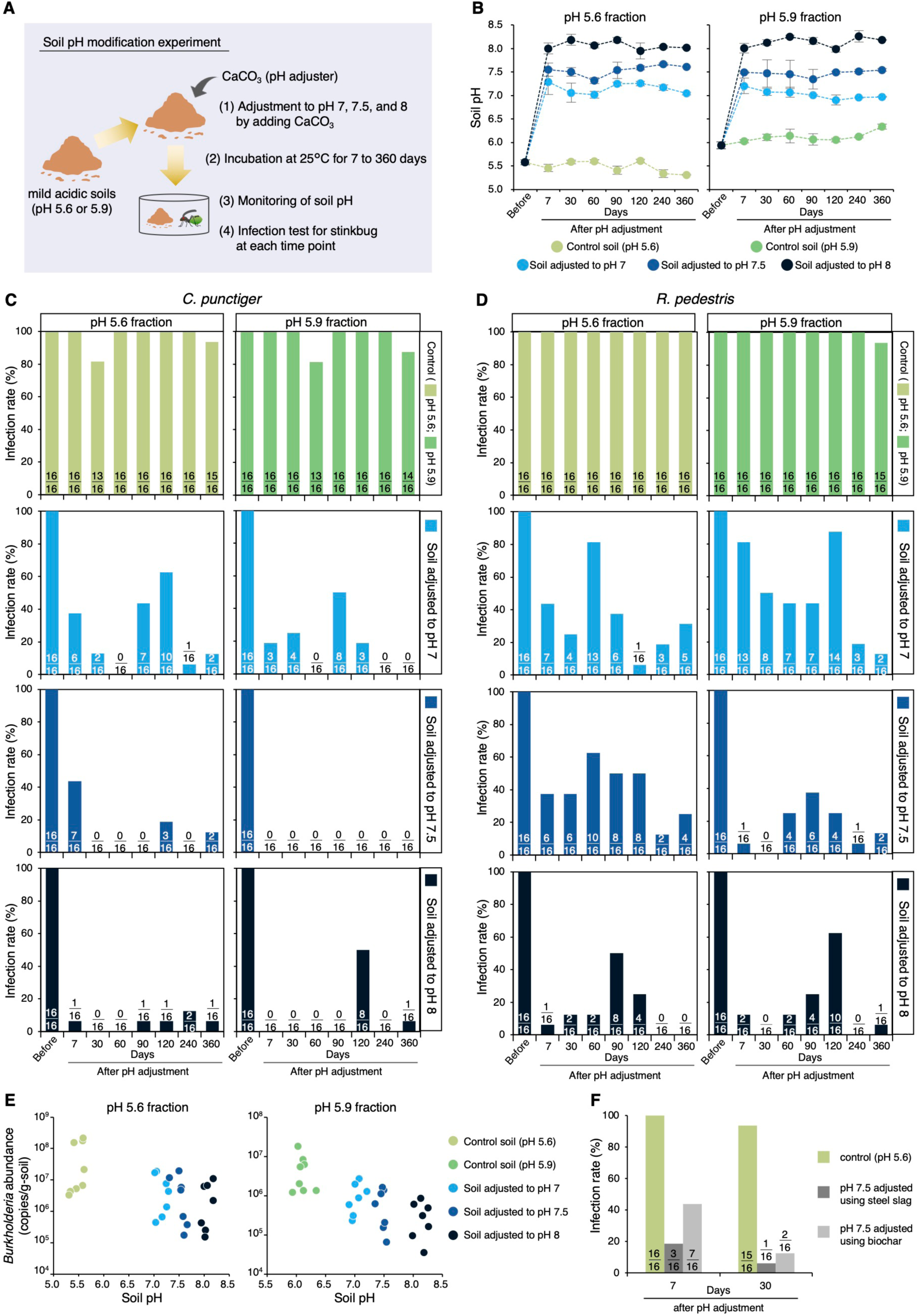
Effect of soil pH modification on symbiosis establishment. (A) Schematic representation of the soil pH modification experiment. The mildly acidic soils with pH 5.6 and pH 5.9 correspond to the S32 and S33 soils, respectively (Table S3). (B) Transition in soil pH following modification to 7–8 via the addition of CaCO_3_ (n = 3). Error bars represent standard deviation (n = 3). (C) and (D) Transition in the infection rate of *C. punctiger* (C) and *R. pedestris* (D) after soil pH modifications. Numbers with bars indicate “the number of infected individuals/total number of examined individuals.” (E) Correlation between soil pH and the abundance of *Burkholderia* in the soil during the experiments. (F) Effect of soil pH modification on the infection rate using steel slag and chicken manure biochar. Numbers with bars indicate “the number of infected individuals/total number of examined individuals.”

### Immediate inhibition of symbiosis via pH-induced loss of symbiont motility

Notably, 7 days after soil pH modification, the infection rate of stink bugs with *Burkholderia* decreased drastically (Fig. 4C–D), although no reduction in *Burkholderia* abundance was observed in the soil compared with the pre-adjustment levels (Fig. S6). This result suggests that the establishment of stink bugs–*Burkholderia* symbiosis is not solely dependent on the amount of *Burkholderia* in the soil.

A previous study showed that *Burkholderia* must pass through a narrow constricted region (CR) in the midgut to colonize the symbiotic organ (Fig. 1C and Fig. S1), and this process depends on flagellar motility: *Burkholderia* uses flagellar motility to pass through the CR, but non-motile mutants cannot enter this CR [22]. As bacterial flagella are driven by a proton gradient across the cell membrane, motility is enhanced under mildly acidic conditions (pH <7.0) as seen in *Escherichia coli* and *Salmonella enterica* [35]. *In vitro* observations confirmed that *Burkholderia* cells expressed active motility at pH 5.5–6.5, whereas most cells lost their motility at pH 7.5 or 8.0 (Fig. 5A–B). Direct observations of the flagella showed that most cells retained their flagella at pH 6.0 but were absent at pH 8.0 (Fig. 5C–D). Furthermore, *in vivo* observations using *Burkholderia* cultured at various pH levels showed that, at pH <7.0, *Burkholderia* passed through the CR to the midgut’s fourth section with bulb (M4B) in the insect gut, while those cultured at pH ≥7.0 remained in the midgut’s third section (M3) without passing through the CR (Fig. 5E and Movies S1–S2). These results suggest that symbiosis establishment is immediately inhibited under pH ≥7.0 conditions and that soil pH plays a crucial role in modulating the direct negative impact of soil pH modification on *Burkholderia* motility. Feeding stink bugs with soil suspensions at various pH levels revealed that the pH of the M3 region, located in front of the CR, remained stable at <7.0 (Fig. S7). The flagellar motility of *Burkholderia* was restored *in vitro* when the pH was reverted from ≥7.0 to 6.0 (Fig. S8), suggesting that there is a mechanism in the M3 region that prevents motility recovery, independent of pH. It is known that various antimicrobial substances are secreted in the M3 region [24], and the stressful environment inside the insect gut may hinder the recovery of motility in *Burkholderia* symbionts.

**Figure 5.**
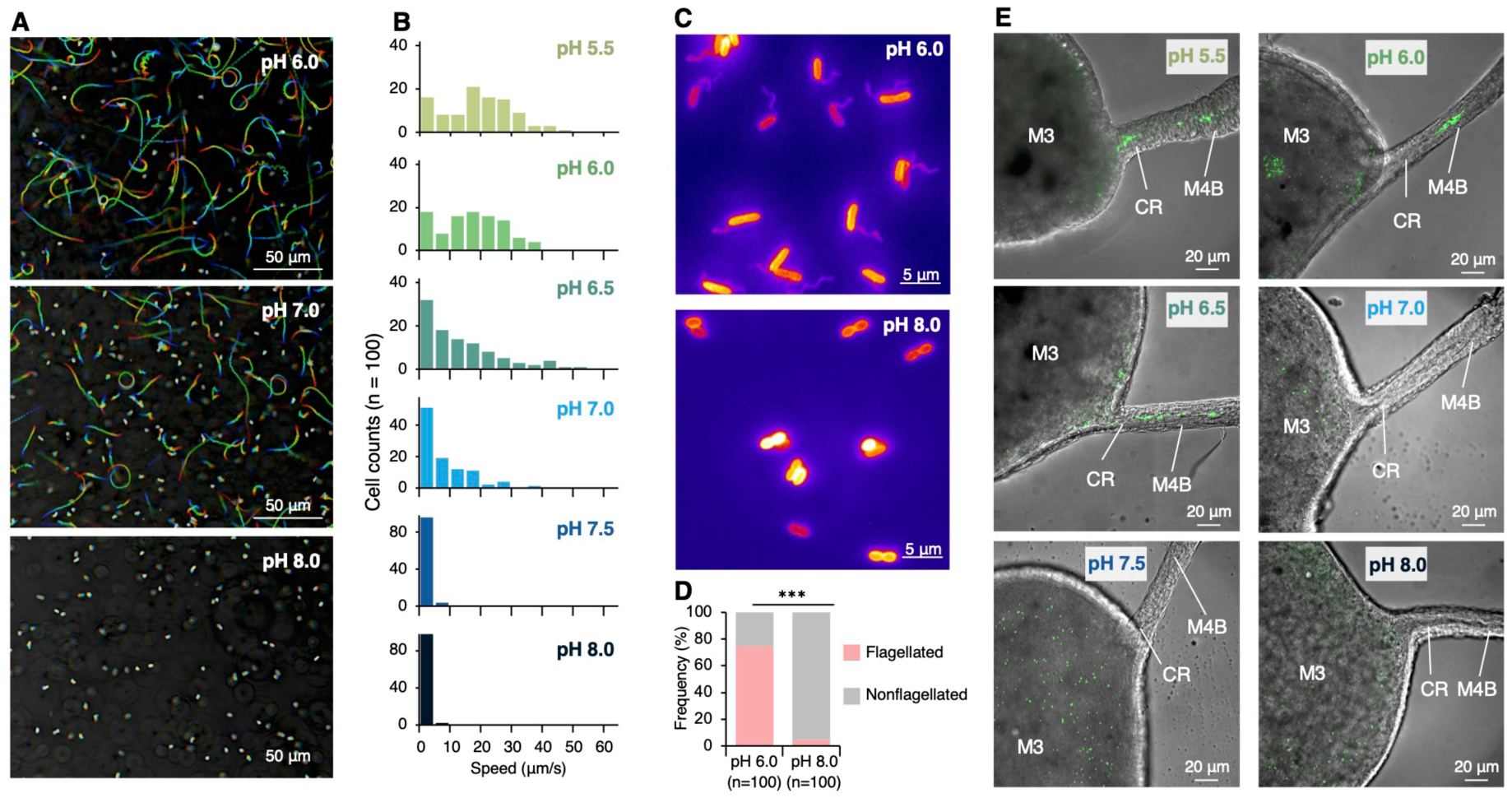
Rapid impact of pH modification on the motility of *Burkholderia* observed through *in vitro* and *in vivo* experiments. (A) Movement trajectories (direction: red to blue) of *Burkholderia* in broth medium with pH 6.0, 7.0, and 8.0 over a 2-s observation period. (B) Swimming speed distribution of *Burkholderia* cells (n = 100) recorded 4 h after pH adjustment. (C) Flagellar observation of *Burkholderia* cells 4 h after pH adjustment using fluorescence microscopy. (D) Proportion of flagellated cells calculated using fluorescence microscopy (n = 100). Statistical analysis was performed using Fisher’s Exact Test: *** (*P* < 0.001). (E) *In vivo* observation; Path (CR) to the symbiotic organ in the midgut of *R. pedestris* fed with *Burkholderia* treated at pH 5.5–8.0. The green signals represent GFP-labeled *Burkholderia*. Abbreviations: M3, midgut third section; CR, constricted region; M4B, midgut fourth section with bulb.

## Discussion

This study demonstrates that soil pH plays a pivotal role in the establishment of symbiosis between stink bugs and their beneficial gut bacteria. Using a multi-scale approach integrating field surveys, soil chemical profiling, laboratory rearing experiments, and bacterial physiology, we found that mildly acidic soils (pH <7.0) facilitate efficient acquisition of *Burkholderia* symbionts, whereas neutral to alkaline soils (pH ≥7.0) disrupt this process (Figs. 3 and 4). This disruption reflects both a reduction in the environmental abundance of the symbiont and a loss of its flagellar motility—an essential trait for host colonization (Figs. 4E and 5). These findings demonstrate that soil physicochemical properties can function as external filters, providing strong evidence that environmental pre-selection of symbiotic bacteria is possible. Although most research has focused on internal partner-choice mechanisms, our results suggest that external environmental selection may play an equally, or even more, important role in shaping the evolution of host–symbiont relationships.

Although the insects studied here harbor environmentally acquired (*i.e*., horizontally transmitted) symbioses, many terrestrial invertebrates possess vertically transmitted symbioses, where the symbionts are inherited from the parental individuals via diverse manner such as coprophagy and transovarial transmission [36]. Such vertically transmitted systems are notably rare in marine animals but relatively common in terrestrial invertebrates [37]. Our findings highlight a fundamental vulnerability of environmentally acquired symbioses in terrestrial habitats: when soil pH exceeds 7.0, stink bugs fail to acquire their beneficial partners. We propose that this ecological uncertainty may have acted as a selective pressure favoring the evolution of vertical transmission in insects. The concept of “partner assurance” has been hypothesized as a major advantage of vertical transmission [38], and our results provide empirical support for this idea in the context of soil-mediated symbioses. In this context, it is noteworthy that *C. schmidti* examined in this study (Table S1), squash bug *Anasa tristis* (Coreoidea), and oriental chinch bug *Cavelerius saccharivorus* (Lygaeoidea) exhibit partial vertical transmission of *Burkholderia* symbionts [39, 40], potentially representing intermediate stages in the evolutionary transition from environmental acquisition to vertical inheritance.

Because many symbiotic microorganisms are essential for their hosts, the distribution of microbes in the environment may directly affect not only insect population dynamics but also their geographic distribution. While insect population dynamics are conventionally attributed to factors such as food availability, natural enemies, or climatic conditions [41–44], our findings reveal that soil chemical properties—particularly pH—can directly determine the success or failure of symbiosis establishment (Fig. 6), which in turn has profound consequences for host growth and reproduction (Figs. 1G and 2). Although previous studies have revealed that soil pH is a major determinant of soil microbial communities [8, 28–33], how this abiotic factor shapes the distribution of terrestrial animals remains poorly understood. Moreover, despite increasing recognition that some animal–microbe associations depend on soil microorganisms [45–47], the role of abiotic soil factors in regulating these interactions has been largely overlooked in both soil science and entomology. By bridging this gap, our study highlights a cross-kingdom effect of soil pH, whereby pH-mediated shifts in microbial communities cascade through symbiotic interactions to influence insect ecology.

**Figure 6.**
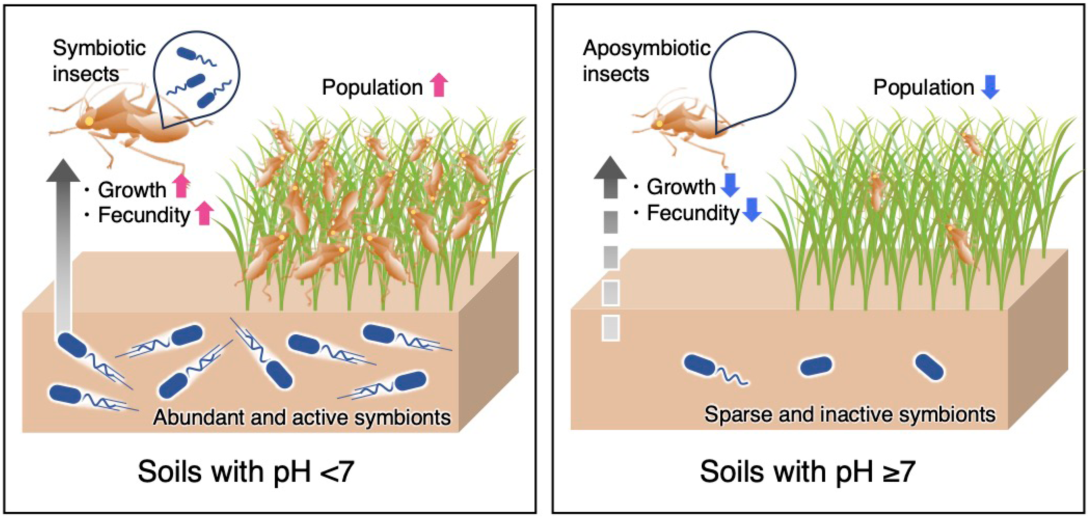
Proposed linkage between soil and insect ecology: Soil pH potentially regulates the ecological success of stink bugs through gut symbiosis. The left and right panels illustrate the interactions among soil pH, soil-derived symbiotic bacteria, and host insects in weedy fields with mildly acidic soils (pH <7.0) and those with neutral to mildly alkaline soils (pH ≥7.0), respectively.

Nevertheless, our field survey did not explicitly account for other ecological variables such as food plant quality, predator abundance, or land management practices including pesticide and herbicide application history. Because our observational data were derived from naturally managed fields, causal relationships between soil pH and insect population dynamics through symbiosis remain to be rigorously confirmed. Future research should therefore employ controlled or semi-controlled environments, such as greenhouse-scale systems, where soil pH and other biotic and abiotic factors can be systematically manipulated. Such experiments will be critical for disentangling the mechanisms by which soil physicochemical properties shape insect–microbe interactions and their broader ecological consequences.

The importance of the gut microbiome is now widely recognized across animal taxa, from vertebrates to invertebrates [48], and there is growing interest in harnessing microbial communities for host control. Most of these strategies focus on manipulating internal microbial communities using prebiotics and probiotics [49]. In contrast, we propose the novel concept of “exobiotics”, referring to environmental interventions that modulate microbial reservoirs outside the host to influence host–microbe interactions. Our findings demonstrate that regulating microbiome dynamics externally can shape the establishment and maintenance of symbiosis. This external dimension has been largely overlooked, except in some marine systems such as corals and bioluminescent squids [6, 50]. By shedding light on this overlooked axis, our work advances a new conceptual framework that emphasizes the triadic relationship among hosts, symbionts, and the environment. This perspective not only enriches symbiosis theory but also provides a scientific foundation for innovative strategies to manage insect populations in agriculture. For example, adjusting soil pH with CaCO_3_, a low-toxicity compound widely used in farming, could hinder symbiont acquisition in weedy breeding grounds of rice stink bugs [51–53], thereby suppressing their population growth and subsequent migration into rice paddies (Fig. S9). The environment has long been regarded merely as a reservoir of symbiotic microorganisms.

This study challenges this view by identifying environmental contexts in which symbionts cannot be acquired, thereby revealing a critical external constraint on symbiosis. These findings highlight the overlooked role of abiotic soil properties in the establishment, maintenance, and evolution of symbiosis, and underscore the importance of symbiont selection processes occurring outside the host.

## Materials and methods

### Insects

We maintained six stink bug species spanning two superfamilies, four families, and five genera: *C. punctiger*, *L. chinensis*, and *P. antennata* collected from weedy fields (mainly *Digitaria ciliaris* and *Setaria viridis*) in Tsukuba (*C. punctiger*) and Ishioka (*L. chinensis* and *P. antennata*), Ibaraki, Japan; *Cletus schmidti* from *Polygonaceae* plants (*Persicaria lapathifolia*) in Sapporo, Hokkaido, Japan; *R. pedestris* from a soybean field in Tsukuba, Ibaraki, Japan [11]; and *M. uniguttatus* from a sparse forest in Naha, Okinawa, Japan (Fig. 1A and Fig. S1). All species except *M. uniguttatus* are agricultural pests [53]. Especially, *C. punctiger* and *L. chinensis* are serious rice pests in Eastern Asia, known to cause sterility and pecky rice [53]. Insects were maintained at 25°C under a long-day regimen (16 h light, 8 h dark), and fed distilled water (DW) and various seeds: *C. punctiger* and *M. uniguttatus* with dry wheat (*Triticum aestivum*) and sunflower (*Helianthus annuus*); *C. schmidti* with wheat, sunflower, and buckwheat (*Fagopyrum esculentum*); *R. pedestris* with soybean (*Glycine max*); *L. chinensis* with wet foxtail millet (*Setaria italica*); and *P. antennata* with wheat, sunflower, southern crabgrass (*Digitaria ciliaris*), green foxtail (*Setaria viridis*), and Japanese millet (*Echinochloa esculenta*). To maintain symbiotic rearing lines, soil S32 (Table S3) was routinely added to DW and provided to hatchlings. Insect samples used in each experiment are listed in Tables S6 and S7.

### Soils

Forty-two soil samples were collected from weedy and agricultural fields across Japan (Table S3). Plant residues and stones were removed with a 2-mm sieve before measuring soil pH and water content. Soil pH was measured by mixing sieved soil with DW (soil/water; 1/5, *w*/*v*) and using a pH meter (LAQUAtwin pH-22B, Horiba, Kyoto, Japan). Water content was determined by weighing several grams of sieved soil before and after drying at 110°C for 15 h. The maximum water-holding capacity (MWHC) of soils S32 and S33 was measured using Hilgard’s method [54]. Other soil chemical properties were measured and analyzed according to standard methods [55]: electric conductivity (EC; platinum electrode method); cation exchange capacity (CEC; semi-micro Schollenberger method); exchangeable calcium and magnesium (atomic absorption spectrometry); exchangeable potassium (flame photometry); available phosphate (Truog method); phosphate absorption coefficient (ammonium phosphate solution method); NH_4_^+^-N (indophenol-blue method); NO_3_^−^-N (UV spectrophotometry); total carbon and nitrogen (dry combustion method); and humus (Kumada method). All measured values were summarized in Table S3. Soil classifications were inferred from collection site coordinates using the Japan Soil Inventory (NARO, https://soil-inventory.rad.naro.go.jp/). Sieved soil was stored at 4°C for the rearing/infection experiments and DNA extraction.

### DNA preparation

The midgut regions of M4B and M4 (symbiotic organs) were dissected from each insect using fine forceps and tweezers under an optical microscope (S9D, Leica Microsystems, Wetzlar, Germany) in a plastic Petri dish filled with phosphate-buffered saline (PBS; 137 mM NaCl, 2.7 mM KCl, 8.1 mM Na_2_HPO_4_, 1.5 mM KH_2_PO_4_, pH 7.5). After digestion in ProK buffer [0.1 M NaCl, 10 mM Tris–HCl (pH 8.0), 1 mM EDTA, 0.2 mg/ml proteinase K] at 56°C for 2 h, the resulting solution was purified by phenol:chloroform:isoamyl alcohol (25:24:1) extraction and ethanol precipitation, as described previously [26]. The quality of the extracted DNA from the insect samples was checked by PCR amplification of the insect cytochrome oxidase I (COI) gene with a universal primer set, LCO1490 and HCO2198 [56] for *C. punctiger*, *C. schmidti*, *L. chinensis*, *R. pedestris*, and *M. uniguttatus*, or the 18S rRNA gene with the primer set, pan18S1552F (5′-CTGTGATGCCCTTAGATGTT-3′) and pan18S1908R (5′-ACCTACGGAAACCTTGTTAC-3′) for *Pachygrontha antennata*.

Soil DNA extraction was conducted by considering 0.25 g (wet weight) of each soil sample using the ISOIL for Beads Beating (Nippon Gene, Tokyo, Japan) according to the manufacturer’s instructions, with the following additional treatments: the addition of 0.02 g skim milk to the lysis buffer to improve DNA extraction efficiency [57] and post-elution purification using ribonuclease A (Nippon Gene). The resulting extracts were further purified using OneStep PCR Inhibitor Removal Kits (Zymo Research, Irvine, CA, USA) to remove the PCR inhibitors as much as possible. The integrity and quantity of the purified DNA were determined using a NanoDrop ND-1000 spectrophotometer (NanoDrop Technologies, Wilmington, DE, USA) and a Qubit 4 Fluorometer (Invitrogen, Carlsbad, CA, USA) with Qubit dsDNA HS Assay Kits (Invitrogen), respectively.

### Diagnostic PCR for *Burkholderia* infection

To determine *Burkholderia* infection in insect samples, diagnostic PCR was conducted using specific primer sets, Bf and Br, for the 16S rRNA gene of *Burkholderia sensu lato* [58]. Previous studies have indicated that stink bugs can infect with *Burkholderia sensu stricto*, including the *B. cepacia* complex and *B. pseudomallei* clade, *Paraburkholderia* (*Burkholderia* spp. in plant-associated beneficial and environmental clade), and *Caballeronia* (*B. glathei* clade or *Burkholderia* spp. in stink bug-associated beneficial and environmental clade) [27, 40, 46], as also shown in this study (Fig. S2). Therefore, this study covers *B. sensu lato*, including these previous *Burkholderia* clades [59]. PCR amplification was performed using an AmpliTaq Gold 360 Master Mix (Invitrogen) according to the manufacturer’s instructions under the following thermocycling conditions: 95°C for 10 min, followed by 25 cycles of 95°C for 30 s, 55°C for 60 s, and 72°C for 30 s. The PCR products were stained with Novel Juice (GeneDireX, Las Vegas City, NV, USA) and checked by electrophoresis on a 2% agarose gel.

### Microbiome analysis of the insect gut and soil

Deep sequencing of the PCR amplicon of the variable region (V4) of the bacterial 16S rRNA gene was performed for DNA samples of soils (Table S3) and insects (Table S7). DNA libraries for the Illumina sequencing platform were prepared and sequenced using 515F and 806R primers [60] on an Illumina iSeq 100 sequencer (Illumina, San Diego, CA, USA), as described previously [18]. Raw sequencing reads were first processed to remove primers, perform quality filtering, and assemble paired-end reads using Trimmomatic ver. 0.39 [61] with the SLIDINGWINDOW:20:30 setting, followed by fastq-join from ea-utils v1.1.2 (https://github.com/ExpressionAnalysis/ea-utils). The resulting assembled sequences were further curated in Mothur ver. 1.42.3 [62]: sequences shorter than 200 bp or longer than 300 bp were discarded using the screen.seqs command, and chimeric sequences were detected and eliminated with the chimera.uchime command using reference-based alignment against the SILVA database (release 128) [63]. Taxonomic classification was conducted with the RDP classifier ver. 2.13 [64], applying a 50% confidence cutoff. Sequences identified as chloroplasts were removed, retaining only those assigned to the kingdom Bacteria for downstream microbiome analyses. All steps were carried out following the procedure described in a previous study [65].

To estimate bacterial abundance, quantitative PCR (qPCR) was performed to amplify the V4 of the bacterial 16S rRNA gene using the LightCycler 96 System (Roche Diagnostics, Basel, Switzerland) with THUNDERBIRD SYBR qPCR Mix (Toyobo, Otsu, Shiga, Japan). The reaction mixture comprised 2× THUNDERBIRD SYBR qPCR Mix, 0.3 μM primers 515F and 806R, and soil DNA samples. The PCR conditions were as follows: initial denaturation at 95°C for 30 s, followed by 45 cycles of 95°C for 15 s, 55°C for 30 s, and 72°C for 30 s. The absolute number of bacterial 16S rRNA gene copies was determined based on a standard curve constructed using 10-fold serial dilutions of the target PCR product of *Burkholderia* sp. SFA1, as described previously [66]. Additionally, *Burkholderia* abundance was calculated by multiplying the bacterial abundance through this qPCR by the relative abundance of *Burkholderia* obtained from the above amplicon sequencing.

### Isolation and inoculation of *Burkholderia* symbionts

*Burkholderia* symbionts were cultured and orally inoculated into early nymphs following the established methods for *R. pedestris* [27]. For *C. punctiger*, *R. pedestris*, *L. chinensis*, and *P. antennata*, we used the strains CPU52, RPE64, LCH90, and PAN135, respectively, which were previously isolated from each insect species [26]. For *C. schmidti*, a *Burkholderia* symbiont strain, designated CSC94, was newly isolated from a wild individual collected in Sapporo, Hokkaido, Japan. This strain was obtained by incubating the crypt contents on 1.5% agar plates containing YG medium (0.5% yeast extract, 0.4% glucose, 0.1% NaCl) at 30°C for 2 days and was identified by sequencing the 16S rRNA gene, as described previously [67]. For *M. uniguttatus*, we were unable to culture the *Burkholderia* symbiont from wild individuals and thus used the strain THE68, previously isolated from the gut of *Togo hemipterus*, within the same family (*Rhyparochromidae*) as *M. uniguttatus* [26]. The symbiont strains were precultured overnight in YG medium with shaking at 180 rpm and 30°C. After an additional 2-h culture in fresh YG medium, symbiont cells were collected by centrifugation, suspended in DW adjusted to 10^4^ CFU/mL, and then fed to hatchlings of each insect by soaking the cell suspension in cotton pads.

Once the nymphs molted to the 3rd instar stage, the symbiont-containing water was replaced by symbiont-free DW, and the nymphs were reared to adulthood.

### Detection of the transmission mode

To elucidate the transmission mode of the *Burkholderia* symbiont in stink bug species, a series of rearing experiments were conducted as follows. The symbiont-infection status of the parents was confirmed by diagnostic PCR for the *Burkholderia* symbiont as described above.

i. *Wild population inspection.* To check the prevalence of the *Burkholderia* symbiont in wild populations of stink bugs, adult insect samples collected from fields in Japan (Table S6) were subjected to diagnostic PCR. The data for *R. pedestris* and *L. chinensis* were obtained from a previously published study [68].
ii. *Egg inspection.* The possibility of vertical symbiont transmission via eggs, such as egg-smearing and transovarial manners, was investigated by collecting eggs from wild individuals and performing diagnostic PCR. The data for *R. pedestris* was obtained from a previously published study [11].
iii. *Rearing of nymphs in clean Petri dishes.* Hatchlings from infected parents were reared in a clean Petri dish until the 2nd instar (for *M. uniguttatus* and *P. antennata*), 3rd (for *R. pedestris*), or 4th instar (for *C. punctiger*, *C. schmidti*, and *L. chinensis*) and subjected to diagnostic PCR.
iv. *Rearing of nymphs in clean Petri dishes with soil.* The possibility of environmental symbiont transmission was investigated by rearing nymphs in a clean Petri dish with soil S32 collected from a field in Sapporo, Hokkaido, Japan (Table S3). The soil was suspended in DW and fed to hatchlings during their development, and the resulting insect samples were subjected to diagnostic PCR. Details are described in the “Infection test using soil samples” section of the Materials and Methods.
v. *Rearing of nymphs in clean Petri dishes with infected parents.* To check the possibility of coprophagy, hatchlings were reared in a clean Petri dish with their infected parents with GFP-labeled *Burkholderia* symbionts that were made using the Tn7 minitransposon system, as described previously [69]. After reaching the 2nd instar (for *M. uniguttatus* and *P. antennata*), 3rd (for *R. pedestris*), or 4th instar (for *C. punctiger*, *C. schmidti*, and *L. chinensis*), these insects were subjected to fluorescence microscopy observation or diagnostic PCR.
vi. *Rearing of nymphs in plant pots.* To test the possibility of symbiont transmission from plants, hatchlings of *R. pedestris* were reared in soil pots with soybean plants. The sieved soil S32 was transferred into plastic pots (10 × 8 cm; opening diameter × depth), and a soybean seedling was planted there. After two weeks of growing the soybean plant, 15 hatchlings were released into a pot covered with a transparent tube and a paper lid (Fig. S3A), fed with 5 dry soybeans, and maintained at 25°C under a controlled light cycle of 12 h of light followed by 12 h of darkness. For comparison, to prevent insects from directly contacting the soil, they were also reared in a confined space where the above-ground parts of plants, previously sterilized with 70% ethanol and rinsed with sterilized DW, were covered with non-woven fabric (Fig. S3B). Each experimental pot was prepared with three replicates. After reaching the 3rd instar (10 days of rearing), the collected insects were subjected to diagnostic PCR.

### Fitness measurement

To assess the fitness effects of the *Burkholderia* symbiont in stink bugs, 1st instar nymphs were prepared for both symbiont-infected (Sym) and uninfected (Apo) groups. The Sym group was inoculated with cultured symbionts at the 1st to 3rd instar stages while the Apo group remained uninoculated. Nymphs were reared in clean Petri dishes, and the survival rate (adult emergence rate) was recorded for all six stink bug species. For *C. punctiger* and *C. schmidti*, the nymphal period (time to adulthood), body length, body weight, and forewing length were also investigated, as well as fecundity-related properties. Newly emerged adult females and males were kept separately for 2 weeks, and the size of their reproductive organs (one ovary and one testis per individual) was measured using ImageJ ver. 1.54g [70]. Each pair of newly emerged female and male from the same group (Sym or Apo) and emergence day was reared together and checked for mating behavior twice daily for 2 weeks. Egg number was determined by counting all eggs laid during the 2 weeks following the first day of laying. Hatching rate was calculated by counting hatchlings born from the eggs laid during the 2 weeks after the first birth.

### Infection test using soil samples

Each soil sample collected from the weedy and agricultural fields (Table S3) was mixed with DW (soil/water; 1/5, *w*/*v*). The resulting suspension was soaked into a cotton pad, which was then fed to 10–20 hatchlings in each Petri dish. After the insects reached the 2nd (for *M. uniguttatus* and *P. antennata*), 3rd (for *R. pedestris* and *L. chinensis*), or 4th instar (for *C. punctiger* and *C. schmidti*), they were subjected to diagnostic PCR to detect *Burkholderia* infection. The pH-modified soils, as described below, were tested in the same manner.

### Field survey

A field survey of 29 weedy fields was conducted during the summer season, from July 30 to August 4, 2017, in three adjacent prefectures in Japan to minimize the effects of seasonal and geographical variation (Table S2). Survey sites were selected where both southern crabgrass (*D. ciliaris*) and green foxtail (*S. viridis*) coexisted and were dominant, along with a mixture of other Poaceae grasses, as such fields serve as the primary breeding grounds for Coreoidea and Lygaeoidea rice bugs [51–53]. At each site, a 100-m² plot was established using a measuring wheel (MURATEC-KDS Co., Kyoto, Japan). Surface soil samples (∼1 cm depth) were collected from the four corners and the center of each plot, suspended in DW (soil/water; 1/5, *w*/*v*), and the soil pH was measured on-site using a pH meter (LAQUAtwin pH-22B, Horiba). Plots showing large variations in soil pH (standard deviation > 0.5) were excluded from the survey. Within each 100-m² plot, insects were collected by sweeping 100 times using a 42 cm diameter silk net (Shiga Konchu Fukyusha, Tokyo, Japan), and the number of Coreoidea and Lygaeoidea insects was recorded.

### pH preference of the *Burkholderia* strains

The symbiont strains for stink bugs and type strains of *B. sensu lato* were precultured overnight in YG medium with shaking at 180 rpm and 30°C. After an additional 2 h of culture in fresh YG medium, the cultured cells were washed and suspended in DW. The cell suspension was adjusted to a final concentration of optical density at 600 nm (OD_600_) = 0.01 and incubated at 25°C for 36 h in YG medium adjusted to each pH (ranging from pH 5.5–8.0 in increments of 0.5) using 50 mM potassium phosphate buffer. The OD_600_ after incubation was measured using a microplate reader (Infinite F200 PRO; Tecan Japan Co., Kanagawa, Japan).

### Soil pH modification experiment

Soils S32 (pH 5.6) and S33 (pH 5.9) (Table S3) were used in this experiment. Each 15 g of the sieved soils was transferred into plastic tubes (4.5 × 10.4 cm; opening diameter × depth), and CaCO_3_ (Fujifilm Wako Pure Chemical Co., Osaka, Japan) was added to adjust the soil pH to approximately 7, 7.5, and 8. These tubes were covered with aluminum foil and incubated at 25 °C for 360 days, with water added twice a week to maintain the water content at 50% MWHC. Soil samples were collected at 7, 30, 60, 90, 120, 240, and 360 days after CaCO_3_ addition. These samples were used for DNA extraction to check the density of *Burkholderia* in soils and were also suspended in DW (soil/water; 1/5, *w*/*v*) for both pH measurements and infection experiments with *C. punctiger* and *R. pedestris*, as described above. For soils S32 and S33, triplicate tubes were prepared and analyzed individually. For the infection experiments, soil suspensions from the triplicates were pooled and fed to hatchling insects. Control systems were maintained at 50% MWHC without CaCO_3_ addition. Furthermore, soil pH modification experiments were conducted using agricultural materials derived from industrial waste, steel slag [71] and biochar [72], as alternative pH adjusters instead of CaCO_3_. Steel slag (The Sangyo Shinko Co., Tokyo, Japan) and chicken manure biochar (Sowa Recycle Co., Tokyo, Japan) were added to soil S32 to adjust the soil pH to approximately 7.5. Soil samples were collected at 7 and 30 days after incubation under the same conditions as the CaCO_3_-treated soils and were fed to hatchlings of *C. punctiger* for infection experiments.

### *In vitro* and *in vivo* observations of *Burkholderia* symbiont motility

For *in vitro* observation, we used a previously established assay [23] with *B. insecticola* RPE64^T^ (currently classified as *Caballeronia insecticola*), a symbiotic strain for *R. pedestris*, to assess flagellar motility. Strain RPE64^T^ was cultured overnight at 30°C in 1/4 strength YG medium, transferred to fresh medium, and incubated for an additional 5 h. Then, 50 µL of the culture was inoculated into 1 mL of fresh 1/4 strength YG medium adjusted to pH 5.5–8.0 (in 0.5 increments) using 50 mM potassium phosphate buffer. After 4 h of incubation, flagellar motility was observed using a phase-contrast microscope (IX73, Olympus) equipped with a 20× objective lens (UCPLFLN20X, NA 0.70, Olympus), a CMOS camera (DMK 33UX174, Imaging Source), and an optical table (HAX-0806, JVI). Coverslip were glow-discharged by a hydrophilic treatment device (PIB-10, Vacuum Device), and sample chambers were assembled with coverslips and double-sided tapes. Cell suspensions were poured into the chamber, and both ends of the chambers were sealed with nail polish to prevent drying. Flagellar formation at pH 6.0 and 8.0 was also observed under a fluorescence microscope (IX73, Olympus) equipped with a 100× objective lens (UPLSAPO 100×OPH, NA1.4, Olympus), an optical filter set (Cy3-4040C, Semrock), a mercury lamp (U-HGLGPS, Olympus), and a CMOS camera (Zyla 4.2; Andor). Greyscale images were recorded and saved as uncompressed sequential TIF files. Fluorescent labeling followed a previous protocol [23]. Furthermore, cells incubated for 4 h in pH 6.0, 7.0, and 8.0 were transferred to fresh pH 6.0 medium, incubated for another 4 h, and re-evaluated for motility.

For *in vivo* observation, we followed a protocol using *R. pedestris* and GFP-labeled symbionts as described previously [27]. The GFP-labeled strain RPE64^T^ was cultured overnight in YG medium, transferred to fresh medium, and incubated for an additional 4 h. The resulting culture was mixed 1:1 with 100 mM potassium phosphate buffer (pH 5.5–8.0) and incubated at 25°C for 1 h. This mixture was then provided to 2nd instar nymphs via cotton pads. Six hours after administration, the midgut was dissected, placed on a microscope slide, and covered with a coverslip. Samples were observed using a laser scanning confocal microscope (TCS SP8, Leica Microsystems).

### pH measurement of the insect gut

Third instar nymphs in the Apo group of *C. punctiger* and *R. pedestris* were provided with either DW or each soil suspension derived from soils S32 (pH 5.6), S33 (pH5.9), S34 (pH 6.7), S51 (pH7.0), S35 (pH 7.5), S36 (pH 8.0), and S37 (pH 8.6) (Table S3) as drinking water, as described above. After 1 week of rearing, the midgut of the nymphs was dissected in a plastic Petri dish filled with DW and divided into distinct parts: M1, M2, M3, M4B, and M4. Each part, collected from five insects, was homogenized in 100 µl of DW by pipetting, and this procedure was performed in triplicate. The pH of the resulting homogenates was measured using a pH meter (LAQUAtwin pH-22B, Horiba).

### Statistics

Categorical data comparing symbiotic and aposymbiotic insects, as well as flagellated and nonflagellated *Burkholderia* cells, were analyzed using Fisher’s exact test. Continuous data comparing symbiotic and aposymbiotic insects were first assessed for normality using the Shapiro–Wilk test and for variance homogeneity using the *F*-test. Based on these results, the most appropriate statistical method was chosen from four options: Student’s *t*-test, Welch’s *t*-test, the Mann–Whitney *U*-test, and the Brunner–Munzel test. pH preferences for the growth of *Burkholderia* strains were analyzed using Dunnett’s test, comparing values at different pH levels to those at pH 7.

To evaluate the outbreak of stink bugs, we conducted a sensitivity analysis using the following method. First, a generalized linear model (GLM) with a negative binomial distribution was constructed, using pH as the explanatory variable and the number of adult or nymph stink bugs as the dependent variable. Submodels were then created by sequentially removing one data point from the explanatory variable, generating as many submodels as there were data points. Finally, the deviance of all these submodels was plotted to identify, which data point removal significantly improved the goodness of fit.

GLM with Gaussian distribution was used to analyze the effect of the soil’s chemical properties on total bacterial abundance. In this analysis, abundance as a responsive variable was log-transformed. We used GLM with a beta-binomial distribution to analyze the same effect on the relative abundance of *Burkholderia* in the soil microbiome and the infection rate for stink bugs. We used the likelihood ratio of the constructed model to the null model to obtain the *P* value (LR test). All parameters of the constructed model, the test of over-dispersion, and the coefficient of determination (*R*^2^) are summarized in Tables S4 and S5. We obtained the above values using the *glmmTMB* function in the glmmTMB package [73], the *testDispersion* function in the DHARMa package [74], and manual calculations. Because we conducted an analysis of the effect of all chemical properties on the relative abundance of *Burkholderia* in the soil microbiome, we calculated the *Q* value using the Storey procedure as a method of false discovery rate (FDR) adjustment [75]. Using the *qvalue* function in the qvalue package [76], we obtained the proportion of true null *P* values (*π*_0_ = 0.2597403) using the bootstrap method and then calculated the *Q* values. All statistical analyses were performed using R version 4.4.1.

## Supporting information

Mov S1

Mov S2

## Declarations

### Ethics approval and consent to participate

Not applicable.

### Consent for publication

Not applicable.

### Data and code availability

Sequence data obtained from PCR amplicon sequencing of the bacterial 16S rRNA genes have been deposited in the GenBank/ENA/DDBJ databases under accession numbers PRJNA1099441, PRJNA1100222, PRJNA1100244, and PRJNA1100246 for the insects’ gut, field soils, treated S32 soils, and treated S33 soils, respectively. The 16S rRNA gene sequence of strain CSC94, a *Burkholderia* symbiont for *C. schmidti* isolated in this study, has also been deposited in the GenBank/ENA/DDBJ databases under accession number PP767298. The source codes for the bioinformatics and statistical analyses conducted in this study have been deposited on figshare (doi: 10.6084/m9.figshare.28943831).

## Acknowledgements

We thank Aika Sawaguchi, Natsuki Nakamura, Megumi Okamoto, Haruka Ooi, Madoka Miyazaki (AIST), and Hitoshi Abe (Hokkaido University) for maintaining insects in laboratory and supporting experiments, Tomonari Watanabe (National Agriculture and Food Research Organization (NARO)) for valuable advice on breeding of *L. chinensis*, and Kanako Tago, Masahito Hayatsu, Norikuni Oka, Tsuyoshi Yamane, Tsubasa Ohbayashi, Nobuyuki Endo, Nobuo Mizutani (NARO), Atsushi Nagayama, Wakako Kakazu, (Okinawa Prefectural Agricultural Research Center) and anonymous farmers for providing soil samples. We also acknowledge Kawada Laboratory Co., Ltd. (www.kawada-labo.com) for providing chemical analysis services for the soil samples and Enago (www.enago.jp) for the English language review.

## Author contributions

Conceptualization: H.I. and Y.K.; methodology: H.I., H.S., D.N., S.J., and Y.K.; investigation: H.I., H.S., D.N., S.J., and Y.K.; visualization: H.I., H.S., D.N., and S.J.; funding acquisition: H.I. and Y.K.; supervision: H.I. and Y.K.; writing – original draft: H.I.; writing – review & editing: H.I., H.S., D.N., S.J., and Y.K.

