## Supplementary information for "Soil pH as an external filter shaping insect–microbe gut symbiosis"

**Contents:**

Figures S1–S9

Tables S1–S7

Movies S1 and S2

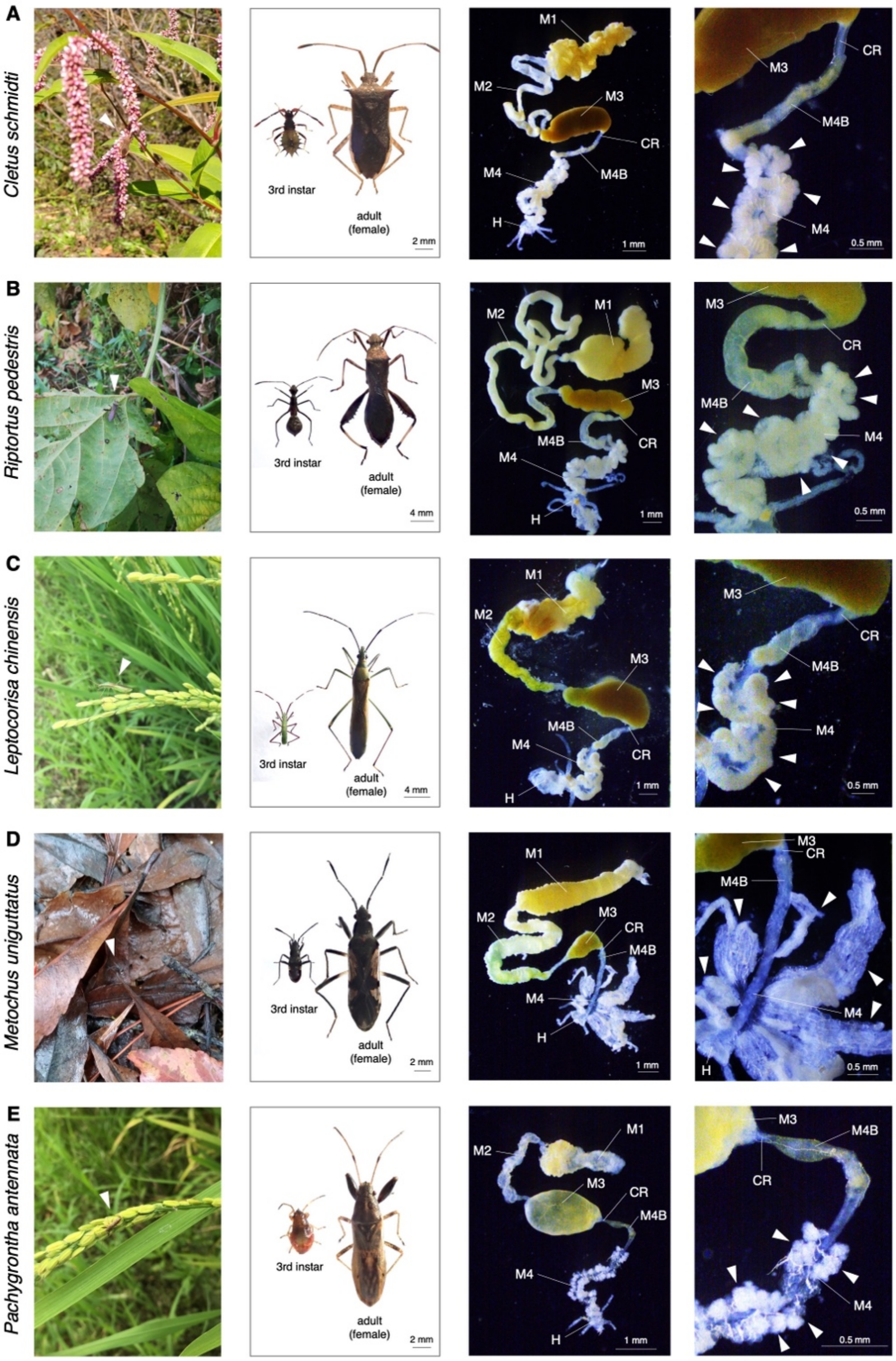


**Fig. S1. Stink bug species used in this study in addition to *C. punctiger*.** The first column displays ecological photographs of each insect species in their natural habitats: (A) *C. schmidti* on the flowers of *Persicaria lapathifolia*; (B) *R.pedestris* on the leaf of *Glycine max*; (C) *L. chinensis* feeding on grains of *Oryza sativa*; (D) *M.uniguttatus* moving on the ground covered with dead leaves; (E) *P. antennata* feeding on grains of *Oryza sativa*. The second, third, and fourth columns depict images of the 3rd instar nymphs and adult females of laboratory-maintained insects, the whole gut, and magnified CR, M4B and M4 regions, respectively. Abbreviations: M1, midgut first section; M2, midgut second section; M3, midgut third section; CR, constricted region; M4B, midgut fourth section with bulb; M4, midgut fourth section with crypts or tubes (symbiotic organ); H, hindgut. Closed triangles indicate crypts and tubes in M4.

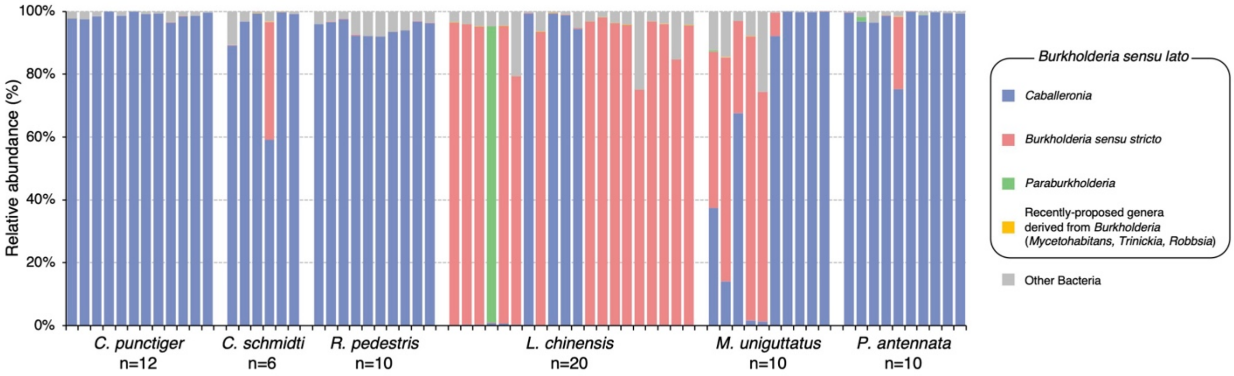


**Fig. S2. Community structure of the gut microbiome in wild populations of stink bug species, estimated by PCR amplicon sequencing analysis of the bacterial 16S rRNA gene.** Detailed information on the insect samples is provided in Table S7. *Caballeronia, Paraburkholderia*, *Mycetohabitans*, *Trinickia*, and *Robbsia* are derivative genera of *Burkholderia* and collectively referred to as *Burkholderia sensu lato* (Mullins and Mahenthiralingam, 2021).

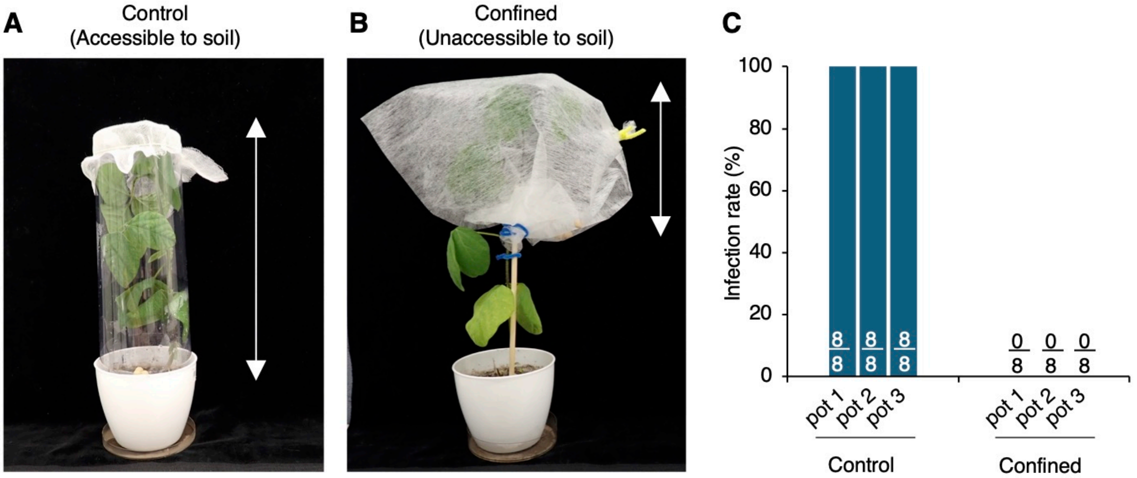


**Fig. S3. Infection experiment conducted in the presence of plants.** (A) and (B) Soybean pots used for the experiment. Soybean seedlings were grown in pots with soil S32. The bidirectional arrow indicates the movable range of insects (*R. pedestris*), showing that insects can access the soil in pots under control conditions (A) but cannot under confined conditions (B). (C) Infection rate of insects with *Burkholderia* after 10 days of rearing in each pot. The experiments were conducted in triplicate pots. Numbers on the bars indicate “number of positive insect samples/total number of examined insect samples.”

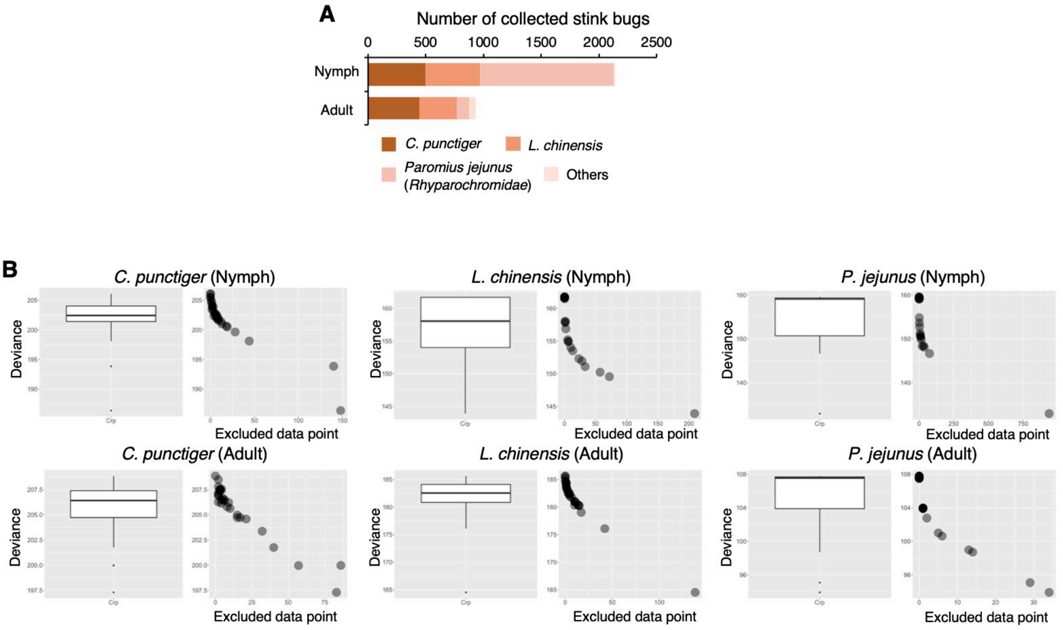


**Fig. S4. Summary and statistical analysis of the field study.** (A) Species composition of all collected stink bug nymphs and adults. Detailed results are provided in Table S2. (B) Sensitivity analysis of stink bug abundance in relation to soil pH. A GLM with a negative binomial distribution evaluated the relationship between soil pH and the number of stink bug species. Submodels excluding one data point at a time identified influential points by comparing the model deviance.

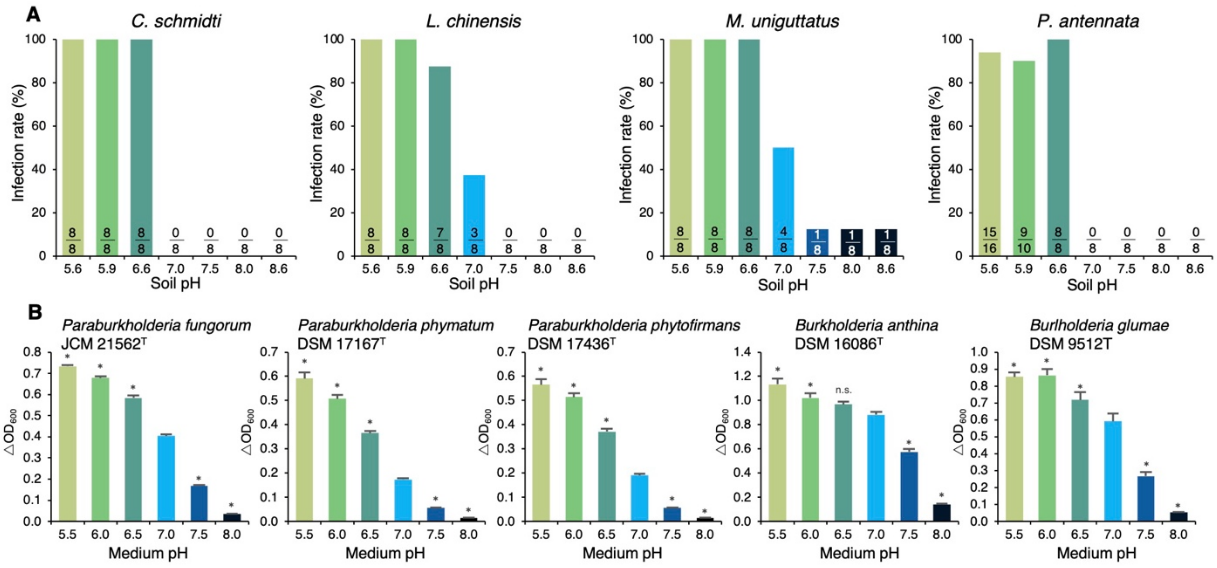


**Fig. S5. pH preferences for symbiosis establishment in various stink bug species and growth of type strains within *Burkholderia sensu lato*.** (A) Infection experiments were performed using seven soils with pH ranging from 5.6 to 8.6, S32 (pH 5.6), S33 (pH5.9), S34 (pH 6.6), S51 (pH7.0), S35 (pH 7.5), S36 (pH 8.0), and S37 (pH 8.6) (Table S3), and four stink bug species: *C. schmidti*, *L. chinensis*, *M. uniguttatus*, and *P. antennata*. Numbers on bars represent "the number of positive insect samples/total number of examined insect samples." (B) pH preferences of three *Paraburkholderia* type strains and two *Burkholderia sensu stricto* type strains were checked using media with pH ranging from 5.5 to 8.0. Error bars represent standard deviation (n = 3). Values marked with an asterisk indicate a significant difference compared to those at pH 7.0, as determined by Dunnett's test: * ($P < 0.05$); n.s., $P \geq 0.05$, not significant.

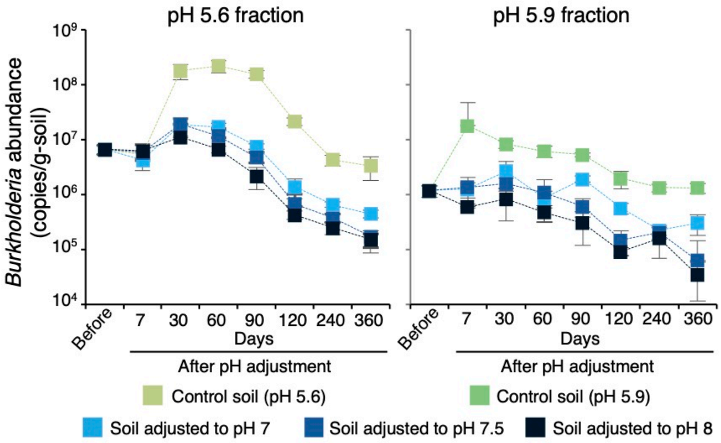


**Fig. S6. Transition of *Burkholderia* abundance in soils after pH modification to levels of 7–8 by $CaCO_3$ addition.** Error bars represent standard deviation (n = 3).

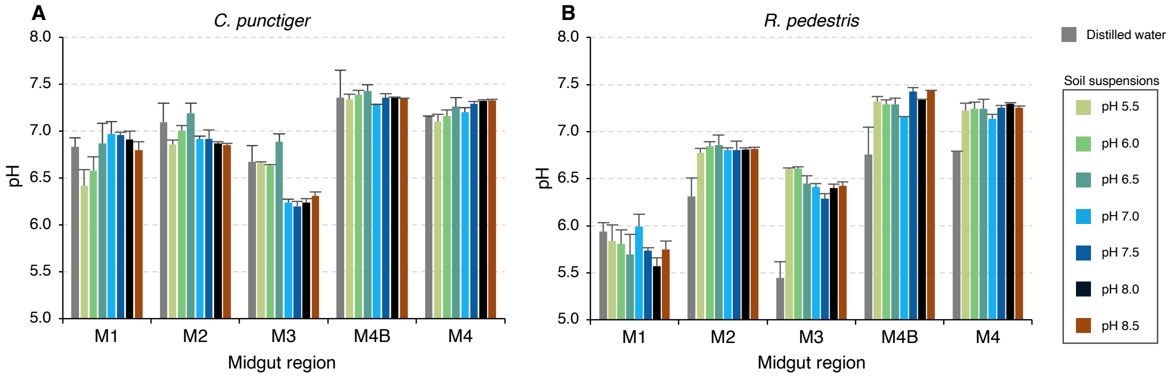


**Fig. S7. Gut pH distribution in stink bugs following the oral administration of soil suspensions with various pH levels.** One week after the oral administration of distilled water or soil suspensions, the midgut sections of the insects were dissected and collected in distilled water for pH measurement (n = 3). (A) and (B) Data from *C. punctiger* and *R. pedestris*, respectively. Abbreviations: M1, midgut first section; M2, midgut second section; M3, midgut third section; M4B, midgut fourth section with bulb; M4, midgut fourth section with crypts (symbiotic organ); H, hindgut. Error bars represent standard deviation (n = 3).

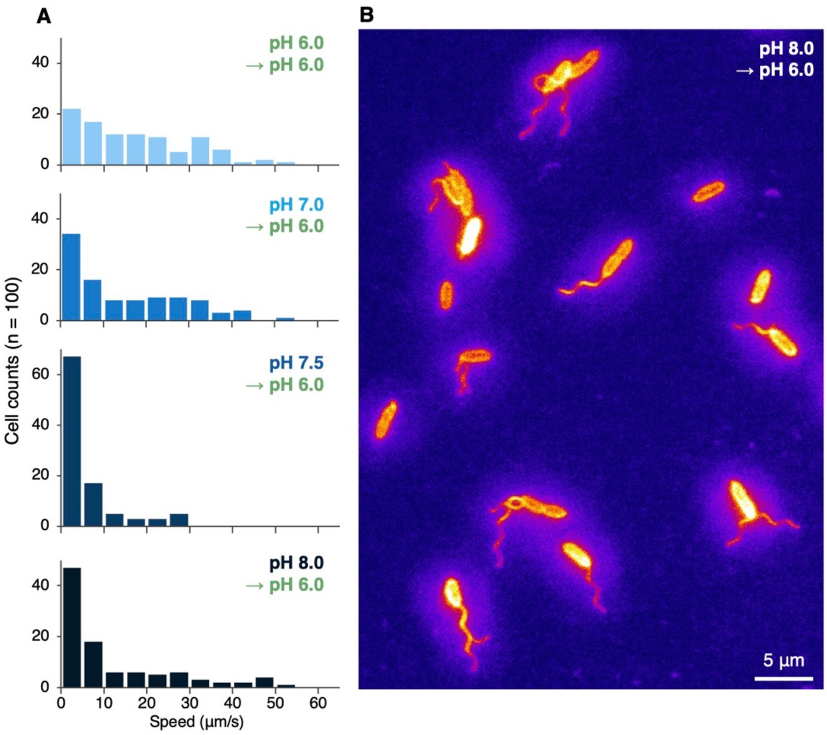


**Fig. S8. *In vitro* observation of flagellar motility in symbiotic *Burkholderia* following pH modifications from 7.0 to 8.0 and then back to 6.0.** (A) Swimming speed distribution of *Burkholderia* cells (n = 100) 4 h after each pH modification. (B) Fluorescence microscopy images of flagella in *Burkholderia* cells captured 4 h after pH modification from 8.0 to 6.0.

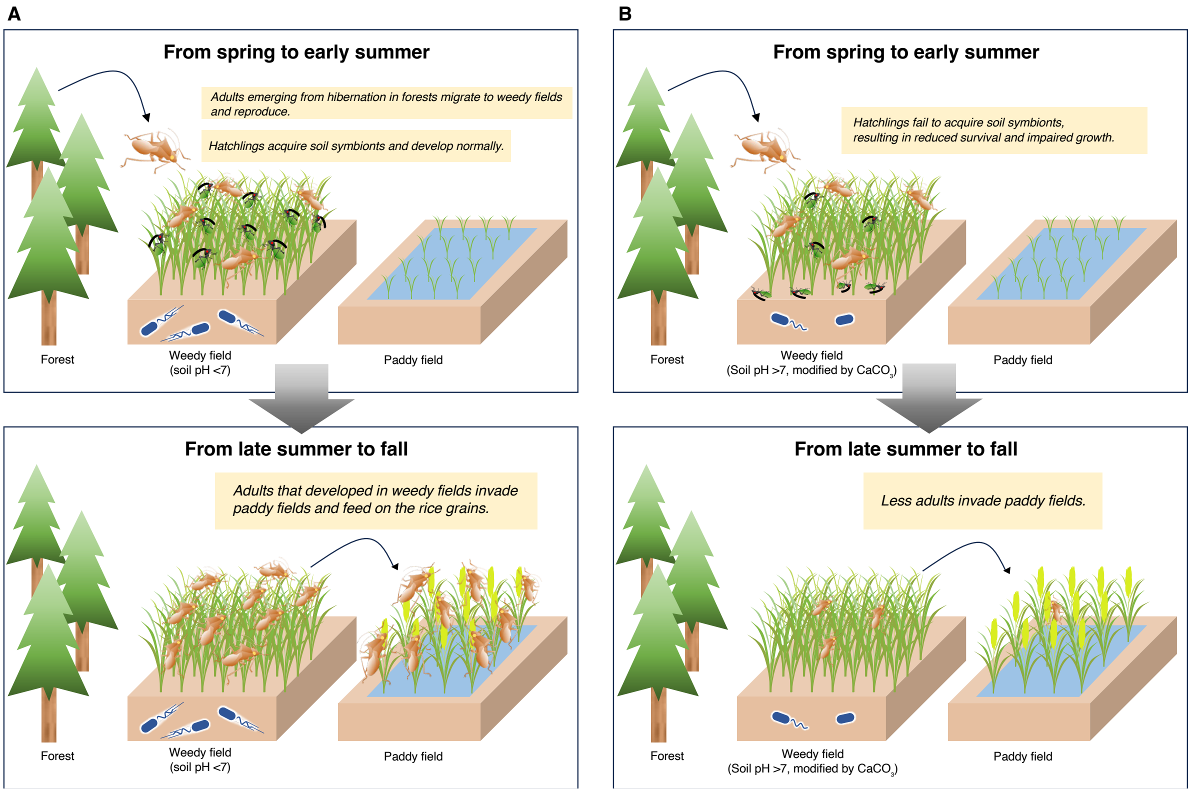


**Fig. S9 Life cycle and invasion pathway of rice stink bugs, and the potential effect of soil pH modification in weedy breeding grounds.** (A) Schematic illustration showing the seasonal life cycle and invasion pathway of rice stink bugs. In spring, adults emerge from hibernation in forests and migrate to weedy fields. From spring to early summer, they reproduce in weedy fields, and the hatchlings develop normally by acquiring symbionts from the soil. Upon reaching adulthood and developing wings, the adults migrate into rice paddies in late summer to autumn, when rice plants enter the grain-filling stage, and feed on the rice grains. (B) Conceptual illustration of the effect of soil pH modification in weedy fields (soil pH >7 adjusted with $CaCO_3$). Even if overwintered adults lay eggs in these fields, hatched nymphs fail to acquire symbionts from the soil, resulting in reduced survival and impaired growth. Consequently, the number of adults invading rice paddies is reduced.

**Table S1. Prevalence of *Burkholderia* among insect samples.**

| Insect taxa | | | Infection rate to *Burkholderia*[a] | | | | |
|---|---|---|---|---|---|---|---|
| Superfamily | Family | Genus Species | Wild population | Eggs | Instars reared without soil and adults | Instars reared with soil[b] | Instars reared with adults |
| Coreoidea | Coreoidae | *Cletus punctiger* | 100% (47/47) | 0% (0/32) | 0% (31/31) | 100% (32/32) | 0% (0/30) |
| | | *Cletus schmidti* | 100% (36/36) | 0% (0/30) | 0% (32/32) | 100% (32/32) | 51.7% (31/60) |
| | Alydidae | *Riptortus pedestris* | 97.6% (87/89)[c] | 0% (0/198)[d] | 0% (0/48) | 100% (8/8) | 0% (0/30) |
| | | *Leptocorisa chinensis* | 98.3% (114/116)[c] | 0% (0/32) | 0% (0/27) | 100% (8/8) | 0% (0/32) |
| Lygaeoidea | Rhyparochromidae | *Metochus uniguttatus* | 100% (10/10) | 0% (0/24) | 0% (0/16) | 100% (8/8) | 0% (0/30) |
| | Pachygronthidae | *Pachygrontha antennata* | 100% (32/32) | 0% (0/32) | 0% (0/16) | 93.8% (15/16) | 0% (0/16) |

[a] Values indicate "the number of positive insect samples/total number of examined insect samples."
[b] Reared with soil S32 (Table S3).
[c] Data from the previous study (Kikuchi et al., 2005).
[d] Data from the previous study (Kikuchi et al., 2007).

**Table S2. Weedy fields surveyed for soil pH and insect density in this study (provided in a separate Excel sheet).**

**Table S3. Soil samples used in laboratory experiments (provided in a separate Excel sheet).**

**Table S4. Results of statistical analyses using Generalized Linear Model (GLM).**

| Constructed model | Factor | Estimate | 95% CI | *P* value | $R^2$ |
|---|---|---|---|---|---|
| **Total bacteria vs pH** | | | | | |
| GLM with Gaussian distribution | Intercept | 10.084 | [9.460, 10.707] | < 0.001 | |
| | pH | −0.062 | [−0.158, 0.034] | 0.205 | |
| Test of dispersion | | | | 0.84 | |
| Coefficient of determination | | | | | 0.075 |
| **Relative abundance of Burkholderia vs pH** | | | | | |
| GLM with betabinomial distribution | Intercept | −2.929 | [−4.578, −1.280] | < 0.001 | |
| | pH | −0.677 | [−0.950, −0.403] | < 0.001 | |
| Test of dispersion | | | | 0.224 | |
| Coefficient of determination | | | | | 0.016 |
| ***C. punctiger* vs pH** | | | | | |
| GLM with betabinomial distribution | Intercept | 21.715 | [12.796, 30.634] | < 0.001 | |
| | pH | −3.223 | [−4.577, −1.869] | < 0.001 | |
| Test of dispersion | | | | 0.768 | |
| Coefficient of determination | | | | | 0.366 |
| ***R. pedestris* vs pH** | | | | | |
| GLM with betabinomial distribution | Intercept | 17.64 | [11.974, 23.307] | < 0.001 | |
| | pH | −2.590 | [−3.441, −1.739] | < 0.001 | |
| Test of dispersion | | | | 0.576 | |
| Coefficient of determination | | | | | 0.312 |
| **Infecion ratio of *C. punctiger* in pH 5.6 fraction** | | | | | |
| GLM with betabinomial distribution | Intercept | 13.885 | [9.320, 18.451] | < 0.001 | |
| | pH | −2.590 | [−2.727, −1.422] | < 0.001 | |
| Test of dispersion | | | | 0.464 | |
| Coefficient of determination | | | | | 0.248 |
| **Infecion ratio of *C. punctiger* in pH 5.9 fraction** | | | | | |
| GLM with betabinomial distribution | Intercept | 23.648 | [12.858, 34.438] | < 0.001 | |
| | pH | −3.531 | [−5.124, −1.939] | < 0.001 | |
| Test of dispersion | | | | 0.744 | |
| Coefficient of determination | | | | | 0.261 |
| **Infecion ratio of *R. pedestris* in pH 5.6 fraction** | | | | | |
| GLM with betabinomial distribution | Intercept | 16.071 | [10.551, 21.590] | < 0.001 | |
| | pH | −2.240 | [−2.988, −1.492] | < 0.001 | |
| Test of dispersion | | | | 0.672 | |
| Coefficient of determination | | | | | 0.304 |
| **Infecion ratio of *R. pedestris* in pH 5.9 fraction** | | | | | |
| GLM with betabinomial distribution | Intercept | 16.709 | [11.020, 22.399] | < 0.001 | |
| | pH | −3.531 | [−3.093, −1.526] | < 0.001 | |
| Test of dispersion | | | | 0.624 | |
| Coefficient of determination | | | | | 0.228 |
| **Burkholderia abundance in pH 5.6 fraction** | | | | | |
| GLM with Gaussian distribution | Intercept | 10.039 | [8.113, 11.966] | < 0.001 | |
| | pH | −2.240 | [−0.755, −0.215] | < 0.001 | |
| Test of dispersion | | | | 0.84 | |
| Coefficient of determination | | | | | 0.145 |
| **Burkholderia abundance in pH 5.9 fraction** | | | | | |
| GLM with Gaussian distribution | Intercept | 10.335 | [8.750, 11.920] | < 0.001 | |
| | pH | −3.531 | [−0.839, −0.401] | < 0.001 | |
| Test of dispersion | | | | 0.84 | |
| Coefficient of determination | | | | | 0.385 |

**Table S5. Correlation between the soil chemical properties and the relative abundance of *Burkholderia* in the soil microbiome, for 42 soils used for the infection experiment (Table S1).**

| Soil chemical properties | *P* value | *Q* value | $R^2$ |
|---|---:|---:|---:|
| pH | **0.000003286** | **0.000011949** | 0.016 |
| Water content (%) | 0.4167 | 0.13261 | 0.001 |
| EC (mS/cm) | **0.006474** | **0.0078** | 0.005 |
| CEC (meq/100 g) | 0.1722 | 0.0895 | 0.001 |
| Exchangeable Ca (mg/100 g) | **0.00002495** | **0.000045364** | 0.013 |
| Exchangeable Mg (mg/100 g) | 0.5677 | 0.15880 | 0.000 |
| Exchangeable K (mg/100 g) | 0.7744 | 0.20114 | 0.000 |
| Available phosphate (mg/100 g) | 0.09977 | 0.06047 | 0.002 |
| Phosphate absorption coefficient (mg/100 g) | **0.03497** | **0.03179** | 0.003 |
| $NH_4$-N (mg/100 g) | 0.2128 | 0.09673 | 0.001 |
| $NO_3$-N (mg/100 g) | 0.09874 | 0.06047 | 0.002 |
| Humus (%) | 0.4364 | 0.13261 | 0.000 |
| Total C (g/kg) | 0.4376 | 0.13261 | 0.000 |
| Total N (g/kg) | 0.4139 | 0.13261 | 0.001 |

**Table S6. Insect samples analyzed for the prevalence of *Burkholderia* in wild populations using diagnostic PCR.**

| Insect species | Collection site | Dominat plant species at the collection site | Collection date | No. of individuals |
|---|---|---|---|---|
| *Cletus punctiger* | Saga, Kanzaki | *Digitaria ciliaris / Setaria viridis* | 29th Oct. 2015 | 19 |
| | Koshi, Kumamoto | *Digitaria ciliaris / Setaria viridis* | 30th Oct. 2014 | 28 |
| *Cletus schmidti* | Takaoka-gun, Kochi | *Achyranthes bidentata* | 1st Nov. 2015 | 12 |
| | Nishiuwa-gun, Ehime | *Achyranthes bidentata* | 1st Nov. 2015 | 24 |
| *Metochus uniguttatus* | Naha, Okinawa | –[a] | 14th Jul. 2019 | 3 |
| | Naha, Okinawa | –[a] | 15th Nov. 2019 | 7 |
| *Pachygrontha antennata* | Ishioka, Ibaraki | *Digitaria ciliaris / Setaria viridis* | 19th Aug. 2020 | 32 |

[a] *M. unigutatus* were captured on fallen leaves in a sparse forest.

**Table S7. Insect samples analyzed for gut microbiota using PCR amplicon sequencing of the bacterial 16S rRNA gene.**

| Insect species | Collection site | Dominant plant species at the collection site | Collection date | No. of individuals |
|---|---|---|---|---|
| *Cletus punctiger* | Tsukubamirai, Ibaraki | *Digitaria ciliaris / Setaria viridis* | 22nd Aug. 2019 | 1 |
| | Kashiwa, Chiba | *Digitaria ciliaris / Setaria viridis* | 22nd Aug. 2019 | 1 |
| | Yaizu, Shizuoka | *Digitaria ciliaris / Setaria viridis* | 2nd Sep. 2019 | 1 |
| | Chiba, Abiko | *Digitaria ciliaris / Setaria viridis* | 22nd Aug. 2019 | 1 |
| | Hamamatsu, Shizuoka | *Digitaria ciliaris / Setaria viridis* | 6th Sep. 2019 | 1 |
| | Yawatahama, Ehime | *Digitaria ciliaris / Setaria viridis* | 2nd Nov. 2015 | 1 |
| | Takaoka-gun, Kochi | *Digitaria ciliaris / Setaria viridis* | 1st Nov. 2015 | 1 |
| | Nakagawa, Fukuoka | *Digitaria ciliaris / Setaria viridis* | 3rd Nov. 2015 | 1 |
| | Yame, Fukuoka | *Digitaria ciliaris / Setaria viridis* | 30th Oct. 2015 | 1 |
| | Oita, Oita | *Digitaria ciliaris / Setaria viridis* | 2nd Nov. 2015 | 1 |
| | Kanzaki, Saga | *Digitaria ciliaris / Setaria viridis* | 29th Oct. 2015 | 1 |
| | Tamana, Kumamoto | *Digitaria ciliaris / Setaria viridis* | 30th Oct. 2015 | 1 |
| *Cletus schmidti* | Sapporo, Hokkaido | *Persicaria lapathifolia* | 2nd Aug. 2014 | 1 |
| | Tsukubamirai, Ibaraki | *Persicaria longiseta* | 23rd Aug. 2019 | 1 |
| | Agawa-gun, Kochi | *Achyranthes bidentata* | 1st Nov. 2015 | 1 |
| | Takaoka-gun, Kochi | *Achyranthes bidentata* | 1st Nov. 2015 | 1 |
| | Nishiuwa-gun, Ehime | *Achyranthes bidentata* | 2nd Nov. 2015 | 1 |
| | Oita, Oita | *Persicaria longiseta* | 2nd Nov. 2015 | 1 |
| *Riptortus pedestris* | Tsukubamirai, Ibaraki | *Glycine max* | 22nd Aug. 2019 | 10 |
| *Leptocorisa chinensis* | Ishioka, Ibaraki | *Digitaria ciliaris / Setaria viridis* | 19th Aug. 2020 | 20 |
| *Metochus uniguttatus* | Naha, Okinawa | –[a] | 14th Jul. 2019 | 3 |
| | Naha, Okinawa | –[a] | 15th Nov. 2019 | 7 |
| *Pachygrontha antennata* | Ishioka, Ibaraki | *Digitaria ciliaris / Setaria viridis* | 19th Aug. 2020 | 10 |

[a] *M. unigutatus* specimens were collected from fallen leaves in a sparse forest.

**Movie S1. The *Burkholderia* treated at pH 6.5, moving within the M4B region in the insect gut (provided in a separate movie file).** The green signals represent GFP-labeled *Burkholderia*.

**Movie S2. The *Burkholderia* treated at pH 7.5, staying within the M3 region in the insect gut (provided in a separate movie file).** The green signals represent GFP-labeled *Burkholderia*.
